# CMG helicase disassembly is essential and driven by two pathways in budding yeast

**DOI:** 10.1101/2024.03.24.586448

**Authors:** Cristian Polo Rivera, Tom D. Deegan, Karim P.M. Labib

## Abstract

The CMG helicase is the metastable core of the eukaryotic replisome and is ubiquitylated and disassembled during DNA replication termination. Fungi and animals use different enzymes to ubiquitylate the Mcm7 subunit of CMG, indicating that CMG ubiquitylation arose repeatedly during eukaryotic evolution. Until now, it was unclear whether cells also have ubiquitin-independent pathways for helicase disassembly and whether CMG disassembly is essential for cell viability. Using reconstituted assays with budding yeast CMG, we generated the *mcm7-10R* allele that compromises ubiquitylation by SCF^Dia2^. *mcm7-10R* delays helicase disassembly *in vivo*, driving genome instability in the next cell cycle. These data indicate that defective CMG ubiquitylation explains the major phenotypes of cells lacking Dia2. Notably, the viability of *mcm7-10R* and *dia2Δ* is dependent upon the related Rrm3 and Pif1 DNA helicases that have orthologues in all eukaryotes. We show that Rrm3 acts during S-phase to disassemble old CMG complexes from the previous cell cycle. These findings indicate that CMG disassembly is essential in yeast cells and suggest that Pif1-family helicases might have mediated CMG disassembly in ancestral eukaryotes.

## Introduction

The duplication of eukaryotic chromosomes initiates at many origins of DNA replication, each of which can only be activated once per cell cycle, to ensure that cells produce a single copy of each chromosome (Bell & Labib, 2016; Costa & Diffley, 2022). The initiation process involves the stepwise assembly of the DNA helicase known as CMG (Cdc45-MCM-GINS), which forms the core of the replisome at replication forks (Fig EV1A). Firstly, two hexameric rings of the six Mcm2-7 ATPases are loaded around double-strand DNA (dsDNA) at origins during G1-phase, to produce inactive Mcm2-7 double hexamers (Evrin *et al*, 2009; Gambus *et al*, 2011; Remus *et al*, 2009). Once cells enter S-phase, Mcm2-7 double hexamers are converted by a set of initiation factors into pairs of CMG helicase complexes that initially encircle dsDNA (Douglas *et al*, 2018; Gambus *et al*, 2006; Lewis *et al*, 2022; Moyer *et al*, 2006; Wasserman *et al*, 2019; Yeeles *et al*, 2015). Subsequently, each pair of CMG complexes is activated in a step that requires the Mcm10 protein and involves extrusion of one of the two parental DNA strands from the Mcm2-7 ring of each helicase (Douglas *et al*., 2018; Lewis *et al*., 2022). This generates two highly stable CMG complexes at the heart of a pair of bidirectional replisomes, with the Mcm2-7 ring of each CMG complex tracking in a 3’ to 5’ direction along single-strand DNA (ssDNA) that represents the template strand for leading-strand synthesis (Fu *et al*, 2011). CMG is essential for DNA replication fork progression and remains continuously associated with the DNA template throughout elongation (Kanemaki *et al*, 2003; Labib *et al*, 2000; Pacek & Walter, 2004; Tercero *et al*, 2000).

When two replication forks from neighbouring origins converge, the CMG helicase is rapidly ubiquitylated and disassembled during DNA replication termination (Dewar *et al*, 2015; Maric *et al*, 2014; Moreno *et al*, 2014). Studies involving *Xenopus* egg extracts (Dewar *et al*., 2015), and a reconstituted budding yeast replisome (Deegan *et al*, 2019; Deegan *et al*, 2020), indicated that the final stretch of parental DNA between the two converging DNA replication forks must be unwound before CMG can be ubiquitylated and disassembled. The converging CMG helicases encircle opposite strands of the parental DNA duplex as they approach, allowing them to bypass each other once the intervening parental DNA has been unwound fully (Low *et al*, 2020). Each helicase then encounters the flapless end of the lagging strand of the opposing fork, allowing both CMG complexes to transition to encircling dsDNA (Fig EV1A). When the two converging CMG helicases bypass each other, this breaks the association of each helicase with the DNA template for lagging strand synthesis that was excluded from the Mcm2-7 ring at replication forks. Throughout elongation, the excluded DNA strand blocks association of the ubiquitin ligase SCF^Dia2^ with a binding site on Mcm2-7 that is essential for ubiquitylation of CMG (Deegan *et al*., 2020; Jenkyn-Bedford *et al*, 2021; Low *et al*., 2020). Thereby, CMG ubiquitylation is restricted to DNA replication termination and represents the final regulated step during the completion of eukaryotic chromosome duplication. SCF^Dia2^ promotes the conjugation of a long K48-linked ubiquitin chain on the Mcm7 subunit of CMG (Deegan *et al*., 2020; Maric *et al*., 2014), leading to recruitment of the Cdc48 ATPase via its ubiquitin receptors Ufd1 and Npl4 (Maric *et al*, 2017). Cdc48 then unfolds ubiquitylated Mcm7 (Deegan *et al*., 2020), triggering disintegration of the CMG helicase and thus inducing replisome disassembly.

Failure to disassemble the metastable CMG helicase during DNA replication termination would leave the Mcm2-7 ring topologically trapped around the DNA duplex, representing a roadblock to a variety of chromatin transactions, including the progression and convergence of DNA replication forks in the subsequent cell cycle. Indeed, budding yeast cells lacking the Dia2 substrate receptor of SCF^Dia2^ are highly sick, have a high rate of genome instability, and are inviable at lower growth temperatures (Blake *et al*, 2006; Koepp *et al*, 2006; Morohashi *et al*, 2009; Pan *et al*, 2006). Structure-guided mutations in the leucine-rich repeats of Dia2, which bind to the Mcm2-7 component of CMG (Jenkyn-Bedford *et al*., 2021), phenocopy the CMG disassembly and cold-sensitivity phenotypes of *dia2Δ* cells (Jenkyn-Bedford *et al*., 2021). However, Dia2 is also likely to use its leucine-rich repeats to bind other substrates, by analogy with other F-box proteins (Harper & Schulman, 2021). Until now, therefore, it was unclear whether the phenotypes of *dia2Δ* cells reflect the consequences of leaving ‘old’ CMG complexes on chromosomes after DNA replication termination, or else result from failure to ubiquitylate other potential substrates of SCF^Dia2^, such as the silencing factor Sir4 (Burgess *et al*, 2012), the transcription factor Tec1 (Bao *et al*, 2004), the licensing factor Cdc6 (Kim *et al*, 2012), the replisome components Ctf4 and Mrc1 (Deegan *et al*., 2020; Mimura *et al*, 2009), or the recombination factor Rad51 (Antoniuk-Majchrzak *et al*, 2023).

In cells lacking Dia2, the CMG helicase persists on chromosomes from DNA replication termination until G1-phase of the next cell cycle (Maric *et al*., 2014). Subsequently, the assembly of new CMG complexes during S-phase is comparable in *dia2Δ* to wild type cells. However, the CMG helicase does not accumulate from one cell cycle to the next in cultures of *dia2Δ* cells and the total amount of CMG is relatively unchanged when *dia2Δ* cells exit G1-phase and progress through S-phase (Maric *et al*., 2014). This suggests that the ‘old’ CMG complexes that persist from the previous cell cycle into G1-phase in *dia2Δ* cells are disassembled during the subsequent S-phase, by a yet unknown pathway (Fig EV1B).

In metazoa, CMG ubiquitylation and disassembly is induced by the ubiquitin ligase CUL2^LRR1^during DNA replication termination (Dewar *et al*, 2017; Fan *et al*, 2021; Le *et al*, 2021; Sonneville *et al*, 2017; Villa *et al*, 2021). Like yeast SCF^Dia2^, CUL2^LRR1^ also ubiquitylates the MCM7 subunit of CMG, in a manner that is repressed at replication forks by the parental DNA strand that is excluded from the Mcm2-7 ring (Jenkyn-Bedford *et al*., 2021), thereby restricting CMG-MCM7 ubiquitylation to termination. However, metazoan CUL2^LRR1^ evolved independently to fungal SCF^Dia2^, and neither enzyme appears to have orthologues in plant genomes. This indicates that CMG ubiquitylation arose more than once during eukaryotic evolution, raising the important question of how CMG disassembly was regulated in ancestral eukaryotes before the evolution of CMG ubiquitylation. Until now, it was unclear whether modern eukaryotes have additional pathways for CMG helicase disassembly that are independent of ubiquitylation and involve factors that evolved before the split between opisthokonts and modern-day plants.

To begin to address all these questions, we developed a mutated allele of Mcm7 that is resistant to ubiquitylation by SCF^Dia2^. Analysis of the new *mcm7* allele indicates that defects in CMG ubiquitylation underlie the major phenotypes of cells lacking Dia2 and lead to genome instability in the subsequent cell cycle. Our data further indicate that the Pif1-family of DNA helicases act during S-phase to mitigate the impact of old CMG helicase complexes from the previous cell cycle, via a second pathway of CMG helicase disassembly that is essential for cell viability in the absence of Mcm7 ubiquitylation. This suggests that Pif1 helicases might have driven an ancient pathway of CMG disassembly in ancestral eukaryotes, predating the evolution of CMG helicase ubiquitylation.

## Results

### Mutation of amino terminal lysines in Mcm7 impairs ubiquitylation and disassembly of the CMG helicase

To explore the consequences of directly impairing CMG ubiquitylation in budding yeast cells, we attempted to develop a form of the CMG helicase that was poorly ubiquitylated by SCF^Dia2^ and Cdc34. This was challenging, since ubiquitin conjugating enzymes (E2 enzymes) do not target specific sites within a consensus sequence, but instead are positioned by ubiquitin ligases (E3 enzymes) in close proximity to their substrates, and thus are able to access a range of potential ubiquitylation sites.

Previously, we showed that the first lysine in budding yeast Mcm7, namely Mcm7-K29 (Fig EV2A-B), is the sole site of *in vitro* CMG ubiquitylation by SCF^Dia2^, when CMG is released from replication fork DNA into budding yeast cell extracts (Maric *et al*., 2017). However, Mcm7-K29 is dispensable *in vivo* for CMG ubiquitylation in budding yeast cells (Maric *et al*., 2017). This suggested that CMG ubiquitylation is more efficient *in vivo* than in yeast cell extracts. Consistent with past findings (Deegan *et al*., 2020), we reconstituted the ubiquitylation of recombinant CMG (Fig EV2C) with purified proteins and found that mutation of Mcm7-K29 blocked CMG ubiquitylation, under conditions where SCF^Dia2^ (E3) and Cdc34 (E2) supported the conjugation of up to about 12 ubiquitins to CMG-Mcm7 (Fig EV2D, lanes 5-6). In contrast, CMG-Mcm7-K29A was robustly ubiquitylated under more efficient conditions that allowed SCF^Dia2^ and Cdc34 to conjugate much longer ubiquitin chains to CMG-Mcm7, though ubiquitylation of CMG-Mcm7-K29A was still less efficient than wild type CMG (Fig EV2D, lanes 7-8). These data showed that SCF^Dia2^ and Cdc34 can ubiquitylate additional sites on CMG-Mcm7 that still remained to be mapped.

Given the failure of mass spectrometry analysis of *in vivo* ubiquitylated CMG-Mcm7 to reveal additional sites of ubiquitylation (Maric *et al*., 2017), we took an alternative approach that was based on the insertion of cleavage sites for the Tobacco Etch Virus (TEV) protease into three poorly conserved disordered loops in yeast Mcm7 (Fig EV2A-B; Fig EV2E; Movie EV1). Ubiquitylation of CMG containing such variants, followed by TEV cleavage, should identify regions of Mcm7 that contain ubiquitylation sites, which could then be characterised further by mutational analysis. Recombinant CMG complexes containing each of the three TEV-site insertions in Mcm7 were purified (Fig EV2F) and shown to be ubiquitylated by SCF^Dia2^ and Cdc34 with comparable efficiency to wild type CMG in reconstituted *in vitro* reactions (Fig EV2G-H). Subsequently, reactions were repeated under the highly efficient conditions shown in Fig EV2D but using lysine-free ubiquitin (K0 Ubi), to block the formation of ubiquitin chains and reveal how many sites were being modified in each molecule of Mcm7 (Fig EV3A). Ubiquitylated CMG complexes were cleaved with TEV protease and the resulting fragments of Mcm7 were then analysed by immunoblotting, with antibodies specific to the amino terminal or carboxy terminal regions of Mcm7 (Fig EV3B-D).

Using CMG with TEV sites inserted after T394 of Mcm7, ubiquitin was conjugated to up to three sites per molecule within the first 394 amino acids of Mcm7 (Fig EV3B, lanes 3-4; note that the assay does not show whether the same three sites are modified in each Mcm7 molecule). In contrast, Mcm7 ubiquitylation downstream of T394 was very inefficient (Fig EV3B, lanes 7-8). These data indicate that SCF^Dia2^ and Cdc34 have a strong preference for ubiquitylating the amino terminal half of Mcm7, consistent with previous work showing that Mcm7-K29 is a favoured site (Maric *et al*., 2017). Subsequently, we analysed cleavage of ubiquitylated CMG complexes with TEV sites inserted after M167 of Mcm7 (Fig EV3C). In this case, up to two ubiquitins were conjugated with high efficiency to the first 167 amino acids, whereas ubiquitylation downstream of M167 was less efficient but still involved up to two ubiquitins per Mcm7 molecule. Combined with the above data, these findings indicated the presence of at least two ubiquitylation sites in Mcm7 1-167, plus up to two sites in the remainder of the protein. Finally, we examined the cleavage products of CMG with TEV sites inserted after A219 of Mcm7 (Fig EV3D) and saw that 2-3 ubiquitins were conjugated to the amino terminal fragment after TEV cleavage (Fig EV3D, lanes 3-4), together with inefficient conjugation of a single ubiquitin downstream of the cleavage sites. In summary, these data indicated that the first 219 amino acids of Mcm7 contain the major sites at which the CMG helicase is ubiquitylated by SCF^Dia2^ and Cdc34.

Finally, we tested whether any of the observed ubiquitylation sites in Mcm7 were induced by TEV cleavage, which might increase the access of SCF^Dia2^ and Cdc34 to residues around the cleavage site. After ubiquitylating CMG-Mcm7(A219-TEV) as above, the sample was split in two and the salt concentration increased in one half to stop the ubiquitylation reaction, before cleavage of Mcm7 with TEV (Fig EV3E). Ubiquitylation of the first 219 amino acids of Mcm7 was unaffected by the presence or absence of high salt during TEV cleavage (Fig EV3F, compare lanes 3 and 7), with 2-4 ubiquitins being conjugated to each Mcm7 molecule in both cases. In contrast, a single ubiquitin was conjugated to Mcm7 downstream of A219 in the control sample (Fig EV3G, lane 3), but this was blocked by increasing the salt concentration (Fig EV3G, lane 7). These data confirmed the presence of multiple ubiquitylation sites within the first 219 amino acids of Mcm7 and further indicated that TEV cleavage leads to inefficient and artefactual ubiquitylation downstream of the cleavage site. Ubiquitylation of the amino terminal region of CMG-Mcm7 was restricted to lysine residues, since mutation of all 10 lysines in the first 219 amino acids to arginine (Fig EV2F, Mcm7-10R(A219TEV)) was sufficient to block ubiquitylation of the Mcm7 amino terminal fragment after TEV cleavage (Fig EV3F, lane 4). These lysine residues are located on the surface of Mcm7, in a region likely to be accessible to Cdc34 when bound to the RING domain of SCF^Dia2^ (Appendix Fig S1; Movie EV2). Mutation of ubiquitylation sites in Mcm7 did not lead to enhanced ubiquitylation of Mcm2-6 (Fig EV3H).

Based on these findings, a version of CMG containing Mcm7-10R but lacking TEV cleavage sites was purified and compared in ubiquitylation reactions to wild type CMG (Fig 1A-B). Under conditions where wild type CMG was ubiquitylated robustly (Fig 1C, lanes 2+8), on up to four sites per Mcm7 molecule (Fig 1C, lane 9), the ubiquitylation of CMG-Mcm7-10R was almost undetectable in reactions containing wild type ubiquitin (Fig 1C, lanes 5+11), though reactions containing lysine-free ubiquitin revealed inefficient ubiquitylation of Mcm7-10R (Fig 1C, lane 12; note that lysine-free ubiquitin provides a more sensitive readout as the products do not smear up the gel lane). Similarly, lysine-free ubiquitin revealed inefficient ubiquitylation of the Mcm3 and Mcm4 subunits of CMG as reported previously (Deegan *et al*., 2020; Maric *et al*., 2017; Mukherjee & Labib, 2019), regardless of the presence of the 10R mutations in Mcm7 (Fig1D-E). In summary, these data indicate that the Mcm7-10R allele greatly reduces CMG ubiquitylation without abolishing it completely.

**Figure 1.**
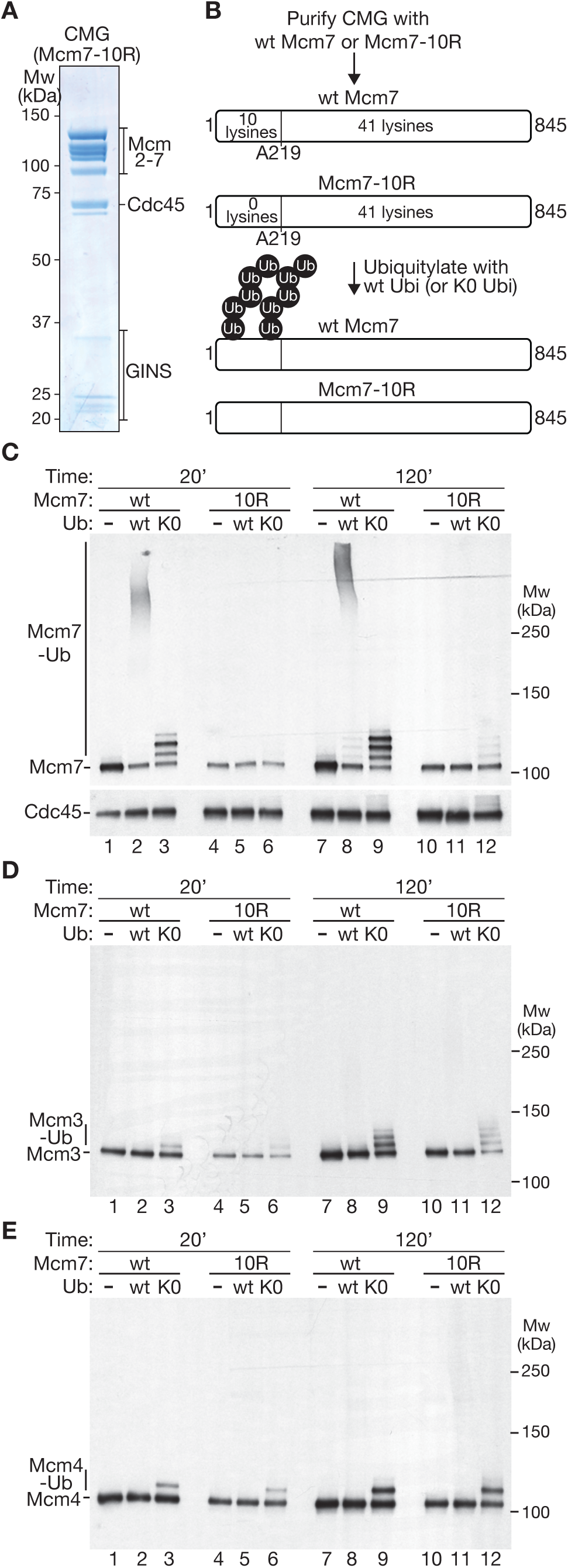
Mcm7-10R impairs *in vitro* ubiquitylation of the CMG helicase by SCF^Dia2^ and Cdc34. (**A**) Purified recombinant CMG helicase with the Mcm7-10R allele. (**B**) Scheme illustrating the ubiquitylation of CMG with wild type Mcm7 (Fig EV2C) or Mcm7-10R (Fig 1A). (**C**) After ubiquitylation of CMG and CMG-Mcm7-10R under the indicated conditions, modification of Mcm7 was monitored by immunoblotting. Cdc45 was included as a loading control. (**D**-**E**) Ubiquitylation of Mcm3 and Mcm4 was monitored in the same experiment as above.

To confirm that the ubiquitylation defect of Mcm7-10R is due to loss of ubiquitylation sites, without affecting the association of CMG-Mcm7-10R with SCF^Dia2^, we incubated wild type CMG or CMG-Mcm7-10R with SCF^Dia2^, in the presence of other replisome proteins that help to link the ubiquitin ligase to the helicase (Deegan *et al*., 2020; Maculins *et al*, 2015; Morohashi *et al*., 2009), and then isolated CMG by immunoprecipitation of the Sld5 subunit of the GINS component (Fig 2A). This showed that SCF^Dia2^ interacted equally well with wild type CMG and CMG-Mcm7-10R (Fig 2B).

**Figure 2.**
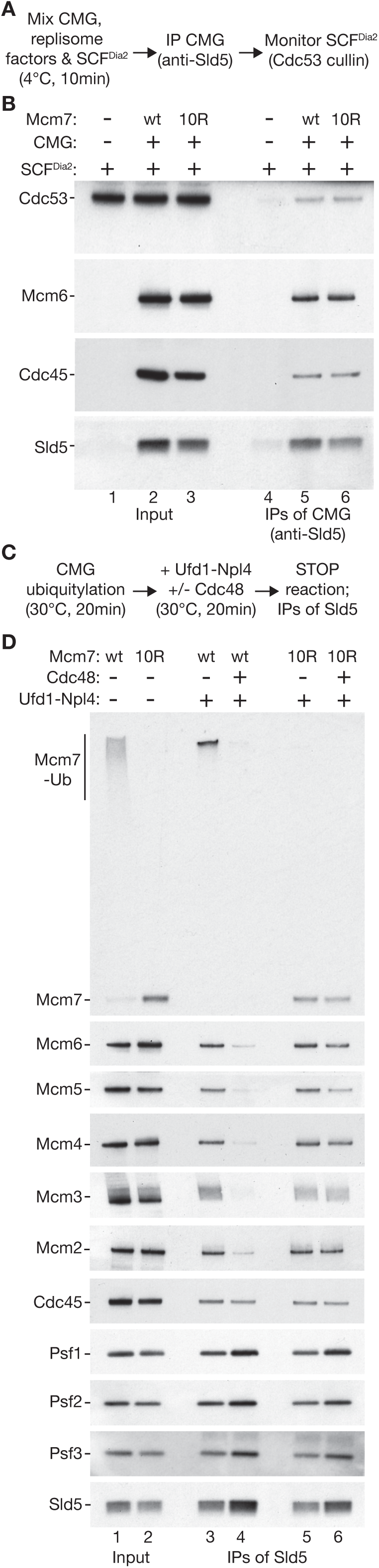
Mcm7-10R associates with SCF^Dia2^ *in vitro* but blocks CMG disassembly. (**A**) Reaction scheme to monitor association of CMG (wt Mcm7 or Mcm7-10R) with SCF^Dia2^ in the presence of the replisome factors Ctf4, Mrc1, DNA polymerase epsilon. (**B**) For the reactions described in (A), the indicated factors were monitored by immunoblotting. (**C**) Scheme for assaying the disassembly of ubiquitylated CMG by Cdc48-Ufd1-Npl4. (**D**) The indicated factors were monitored by immunoblotting, for the reactions described in (C).

To demonstrate that the ubiquitylation defect of Mcm7-10R impairs CMG helicase disassembly, we performed ubiquitylation reactions with wild type CMG and CMG-Mcm7-10R as above. Purified recombinant versions of Cdc48 and its adaptor complex Ufd1-Npl4 were then added to the reaction mixtures for 20 minutes. Subsequently, the reactions were stopped by increasing the salt concentration before incubation with beads coated with antibodies to the Sld5 subunit of GINS (Fig 2C). In reactions containing wild type CMG, addition of Cdc48-Ufd1-Npl4 disrupted the interaction between GINS and the Mcm2-7 complex (Fig 2D, compare lanes 3-4), indicating efficient helicase disassembly as described previously (Deegan *et al*., 2020). In contrast, disassembly of CMG-Mcm7-10R was greatly impaired (Fig 2D, compare lanes 5-6).

### Phenotypic analysis of *mcm7-10R* indicates that CMG is a major target of SCF^Dia2^ in <u>the budding yeast cell cycle</u>

To explore the phenotypic consequences of impairing CMG ubiquitylation and disassembly, the *mcm7-10R* mutations were introduced into the *mcm7* locus in budding yeast cells. Mcm7-10R cells were viable and replicated with similar kinetics to control cells (Fig 3), indicating that the mutation of surface lysines did not impair the assembly or action of the CMG helicase. To examine the impact of *mcm7-10R* on CMG ubiquitylation *in vivo*, we used cells in which Cdc48 was fused to the auxin-inducible degron or ‘AID’ (Maric *et al*., 2014; Nishimura *et al*, 2009), allowing us to degrade Cdc48-AID in early S-phase and thereby monitor the accumulation of ubiquitylated CMG during DNA replication termination. *cdc48-aid* and *cdc48-AID mcm7-10R* cells were released from G1-phase into early S-phase in the presence of 0.2 M hydroxyurea (Fig 3A), to inhibit ribonucleotide reductase and slow the progression of replication forks from early origins, leading to repression of late origin firing via the S-phase checkpoint pathway. Subsequently, auxin was added for one hour to inactivate Cdc48-AID and cells were then released into fresh medium lacking hydroxyurea, allowing them to complete chromosome replication. Samples were taken during early and late S-phase (Fig 3A, samples 1+2) and used to isolate the CMG helicase from cell extracts, by immunoprecipitation of a TAP-tagged version of the Sld5 subunit of GINS. When *cdc48-AID* control cells completed S-phase, CMG accumulated with ubiquitylated Mcm7 and residual ubiquitylation of Mcm3 (Fig 3B, lane 6; note that ubiquitin chains are short under these conditions, likely due to partial depletion of free ubiquitin upon inactivation of Cdc48-A|D). In contrast, ubiquitylation of CMG-Mcm7 was impaired by the *mcm7-10R* mutations, without affecting inefficient Mcm3 ubiquitylation (Fig 3B, lane 8). These data demonstrated that *mcm7-10R* interferes with CMG ubiquitylation in budding yeast cells but does not block it entirely.

**Figure 3.**
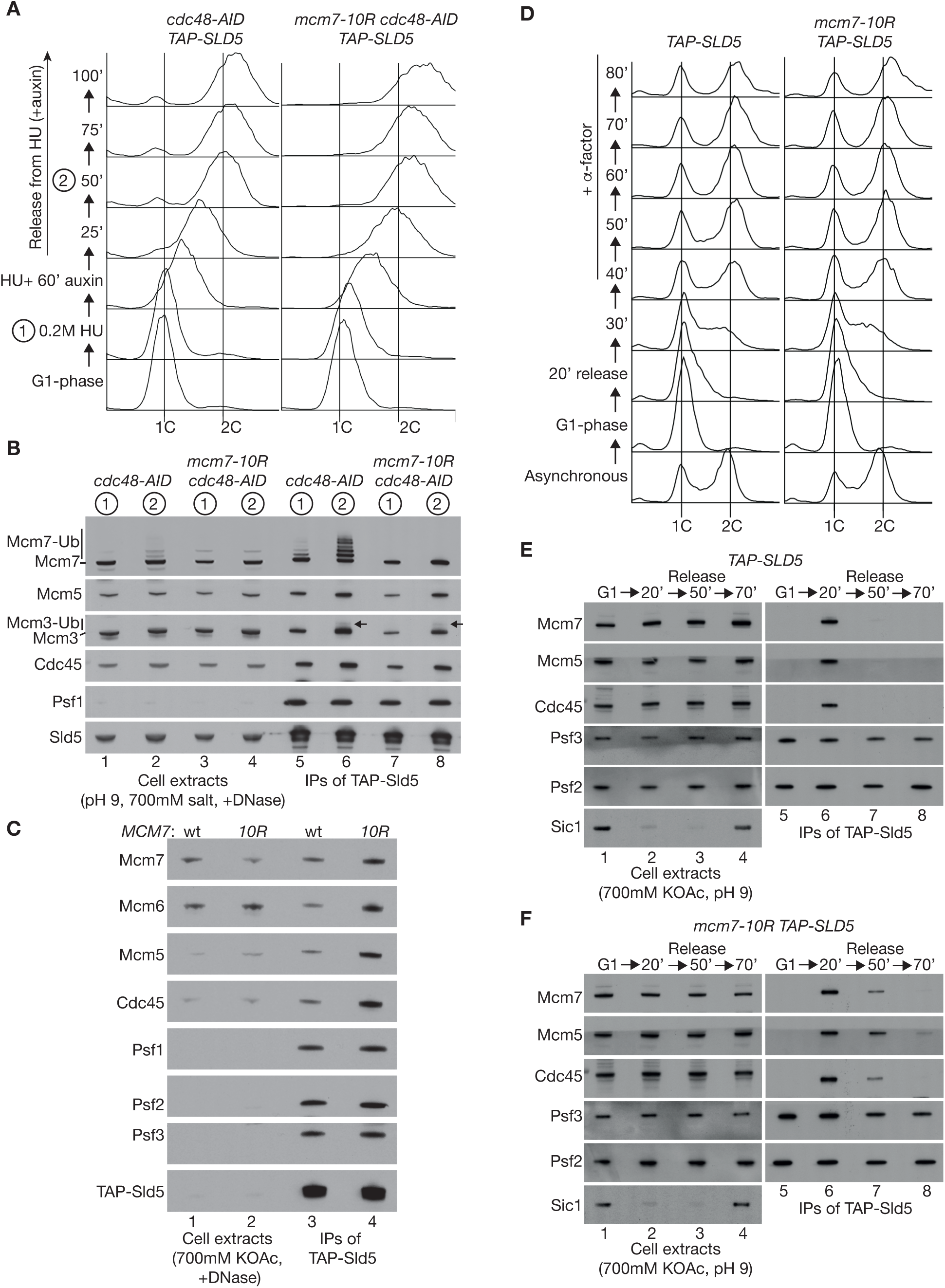
*mcm7-10R* impairs CMG ubiquitylation and delays CMG disassembly in budding yeast cells. (**A**) *cdc48-aid TAP-SLD5* (YPNK337) and *mcm7-10R cdc48-AID TAP-SLD5* (YCPR86) strains were arrested in G1-phase with mating pheromone at 30°C and then released into S-phase for X minutes in the presence of 0.2M hydroxyurea (HU). Subsequently, auxin (0.5 mM IAA) was added for 60 minutes in the continued presence of HU, and cells were then washed into fresh medium containing auxin lacking hydroxyurea, to allow progression through S-phase. DNA content was monitored throughout the experiment by flow cytometry. (**B**) Cell extracts were prepared at the indicated times for the experiment in (A) and used to isolate CMG via the TAP-tagged Sld5 subunit of GINS, under conditions that blocked *in vitro* CMG ubiquitylation. The indicated factors were monitored by immunoblotting. Arrows indicate the small population of ubiquitylated Mcm3 after inactivation of Cdc48-AID. (**C**) TAP-Sld5 was isolated from extracts of asynchronous cultures of control cells (YSS47) and mcm7-10R cells (YCPR79). (**D**) *TAP-SLD5* (YSS47) and *mcm7-10R TAP-SLD5* (YCPR79) strains were arrested in G1-phase with mating pheromone at 30°C and then released into S-phase in fresh medium. Mating pheromone was added again after 40 minutes to prevent cells from entering S-phase in the subsequent cell cycle. (**E**-**F**) TAP-Sld5 was isolated from cell extracts at the indicated time points. Sic1 was also monitored in cell extracts as a marker for arrest in G1-phase.

To monitor the impact of *mcm7-10R* on CMG disassembly, we first examined the level of the helicase in extracts of asynchronous cell cultures, by immunoprecipitation of TAP-tagged Sld5. A similar amount of GINS was isolated from control cells and *mcm7-10R*, but the association of GINS with Cdc45 and Mcm2-7 was enhanced in *mcm7-10R*, indicating accumulation of the CMG helicase (Fig 3C). Subsequently, the assembly and disassembly of CMG were monitored in synchronous cultures of control and *mcm7-10R* cells, following arrest in G1-phase and release into S-phase (Fig 3D). In control cells, CMG was detected 20 minutes after release from G1-arrest, corresponding to cells in early S-phase (Fig 3E, lane 6). Thirty minutes later, CMG was scarcely detectable (Fig 3E, lane 7), reflecting the fact that S-phase takes about 20 minutes in budding yeast. CMG assembly in *mcm7-10R* cells was comparable to the control (Fig 3F, lanes 5-6). However, the disappearance of CMG was markedly slower in *mcm7-10R* cells (Fig 3F, lane 7), indicating that CMG ubiquitylation was impaired but not completely blocked, and suggesting that CMG disassembly under such conditions is driven by residual ubiquitylation of Mcm7, Mcm3 or Mcm4 (c.f. Fig 1C-E).

The impact of *mcm7-10R* on CMG ubiquitylation and disassembly is reminiscent of the effect of deleting the amino terminal TPR domain of DIA2 (TPR = Tetratricopeptide repeat). Previous work showed that the Dia2-TPR domain increases the efficiency of CMG ubiquitylation by binding to Ctf4 and Mrc1 (Deegan *et al*., 2020; Maculins *et al*., 2015), which associate with the CMG helicase in the replisome (Gambus *et al*., 2006). In *dia2-ΔTPR* cells, CMG ubiquitylation is impaired to a lesser degree than in *mcm7-10R*, but the defect is still sufficient to delay CMG helicase disassembly (Maculins *et al*., 2015).

Since CMG disassembly is delayed but not abolished in *mcm7-10R* or *dia2-ΔTPR*, most helicase complexes were disassembled when cells were arrested in G1-phase for an extended period (Fig 4A, Fig 4B lanes 6-7). In contrast, CMG persisted from DNA replication termination to G1-phase of the following cell cycle in *dia2Δ* cells (Fig 4A, Fig 4B lane 5), as shown previously (Maric *et al*., 2014). Notably, the CMG disassembly defects of *mcm7-10R* and *dia2-ΔTPR* are additive, and most CMG complexes persisted into G1-phase of the following cell cycle in the *mcm7-10R dia2-ΔTPR* double mutant (Fig 4A, Fig 4B lane 8), as seen with *dia2Δ*. Moreover, *mcm7-10R dia2-ΔTPR* reproduced other previously reported phenotypes of *dia2Δ* cells (Blake *et al*., 2006; Koepp *et al*., 2006; Morohashi *et al*., 2009; Pan *et al*., 2006), such as cold-sensitivity (Fig 4C), sensitivity to the alkylating agent methyl methanesulfonate (MMS, Fig 4D) and synthetic lethality with *mec1Δ sml1Δ* or *slx8Δ* (Appendix Fig S2). These findings suggested that failure to disassemble the CMG helicase is a major determinant of *dia2Δ* phenotypes. Consistent with this view, mass spectrometry analysis of Protein A-tagged Dia2, isolated from extracts of S-phase yeast cells, indicated that the replisome is the major partner of SCF^Dia2^, largely dependent upon the TPR domain of Dia2 (Appendix Table S1-S2 and Datasets EV1-EV2). In contrast, other reported substrates of SCF^Dia2^ such as Tec1, Rad51, Sir4 and Cdc6 were not either detected or were very weakly enriched in mass spectrometry analysis of immunoprecipitates of Protein A-tagged Dia2 (Appendix Table S1-S2 and Datasets EV1-EV2).

**Figure 4.**
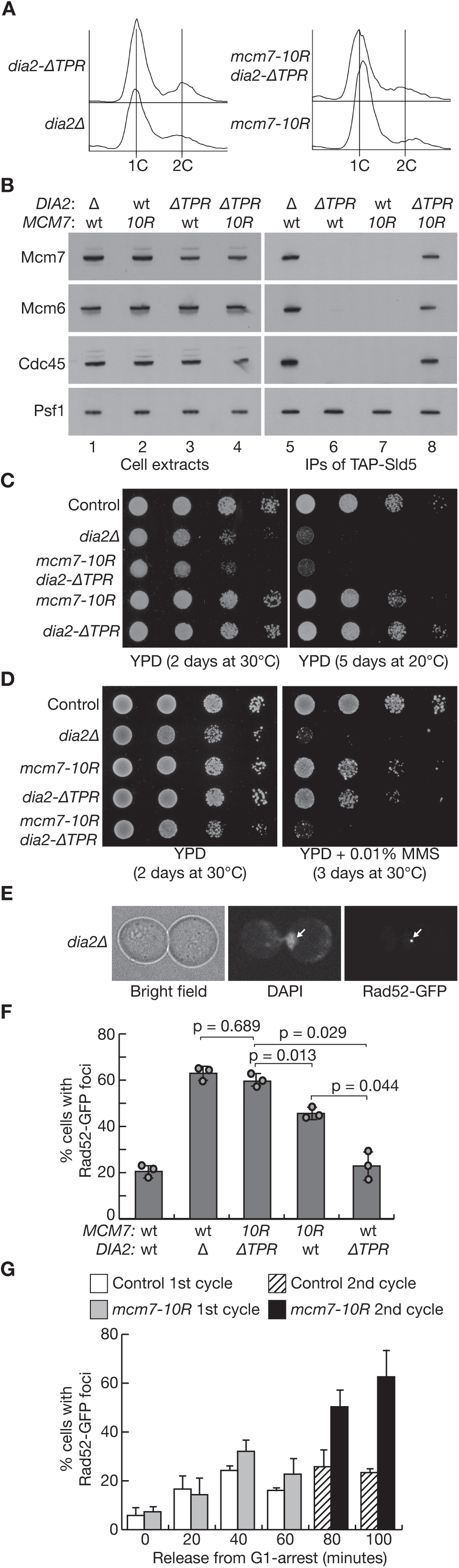
Defects in CMG helicase disassembly phenocopy loss of Dia2 and induce genome instability in the next cell cycle. (**A**) Budding yeast cells of the indicated genotypes (YHM130, YTM265, YCPR86, YCPR111) were arrested in G1-phase at 30°C by addition of mating pheromone. (**B**) GINS and the CMG helicase were isolated from cell extracts by immunoprecipitation of TAP-tagged Sld5. The indicated factors were monitored by immunoblotting. (**C**-**D**) Serial dilutions of the indicated genotypes (YHM28, YHM306, YCPR68, YCPR103 were spotted on media under the indicated conditions. YPD = ‘Yeast extract Peptone Dextrose’ medium. MMS = methyl methanesulfonate. (**E**) Example of *dia2Δ* cell with a sub-nuclear focus on Rad52-GFP. (**F**) The percentage of cells with sub-nuclear foci of Rad52-GFP was monitored for the indicated genotypes (YBH295, YTM115, YCPR447, YCPR445, YTM74). The data from three independent experiments are shown as circles, with the histograms showing mean values and lines indicating standard deviation. Statistical analysis was performed using a one-way ANOVA test followed by Tukey’s test, to generate the indicated p-values. (**G**) Control (YBH295) and *mcm7-10R* (YCPR445) were synchronised in G1-phase by addition of mating pheromone before release into fresh medium lacking mating pheromone. The proportion of cells with sub-nuclear foci of Rad52-GFP was quantified at the indicated timepoints, for cells in the first cell cycle or second cell cycle (the latter were identified as cells that had completed nuclear division and formed a small bud). The histograms indicate the mean values from three independent experiments, with standard deviation indicated by lines.

Cells lacking Dia2 accumulate with sub-nuclear foci of the recombination factor Rad52 (Fig 4E-F), indicating endogenous DNA damage (Blake *et al*., 2006; Morohashi *et al*., 2009). Whereas the proportion of cells with Rad52 foci was not increased significantly in *dia2-ΔTPR* cells, the proportion of *mcm7-10R* cells with Rad52 foci was intermediate between control cells and *dia2Δ* (Fig 4F), reflecting the stronger CMG ubiquitylation defect in *mcm7-10R* compared to *dia2-ΔTPR*. The proportion of cells with Rad52 foci increased further in the *mcm7-10R dia2-ΔTPR* double mutant, to a level that was indistinguishable from *dia2Δ*. Together with the above analysis, and the previous characterisation of structure-guided mutations in the leucine-rich repeats of Dia2 (Jenkyn-Bedford *et al*., 2021), these data indicate that CMG is a major target of SCF^Dia2^ in the budding yeast cell cycle, with the persistence of ‘old’ CMG helicase complexes being a major driver for the phenotypes of cells lacking Dia2.

### Old CMG complexes increase the frequency of Rad52 recombination foci in the next cell cycle

As noted above, the persistence of old CMG complexes in *mcm7-10R* cells can be largely suppressed by arresting cells for an extended period in G1-phase of the subsequent cell cycle, by addition of mating pheromone to the cell culture (Fig 4B, lane 7). This enabled us to compare how defective CMG disassembly drives genome integrity in the first and second cell cycles, by releasing control cells and *mcm7-10R* cells from an extended G1 arrest and then monitoring the kinetics of Rad52 sub-nuclear foci, as cells progressed through the first cell cycle and then entered the second. Cells continued to grow and increase their size during the initial arrest in G1-phase, leading subsequently to rapid entry into S-phase of the first cell cycle in fresh medium lacking mating pheromone, and a very short G1-phase in the second cell cycle.

Upon release into S-phase of the first cell cycle, the proportion of cells with sub-nuclear foci of Rad52-GFP was initially similar in control cells and *mcm7-10R* (Fig 4G, 0-20 minutes), consistent with *mcm7-10R* cells only containing a residual level of old CMG complexes when they entered S-phase. However, when cells completed the first cell cycle and rapidly entered the following cycle, the proportion of cells with sub-nuclear foci of Rad52-GFP was greatly enhanced in *mcm7-10R* cells (Fig 4G, 80-100 minutes; *mcm7-10R* 2^nd^ cycle). These findings indicated that old CMG complexes from one cell cycle represent a source of genome instability when cells enter S-phase of the subsequent cell cycle.

### Pif1-family helicases are essential for viability of *dia2Δ* and *mcm7-10R* and mediate a second pathway for CMG helicase disassembly

When *dia2Δ* cells progress through S-phase, old CMG complexes from the previous cell cycle are removed by an unknown pathway. One candidate factor for such a pathway is the Rrm3 DNA helicase, which together with its paralogue Pif1 is known to help replication forks pass a range of roadblocks on chromatin (Bochman *et al*, 2010; Malone *et al*, 2022; Muellner & Schmidt, 2020), such as tightly bound protein-DNA complexes including Mcm2-7 double hexamers (Hill *et al*, 2020) or stable complexes at tRNA promoters or telomeres (Ivessa *et al*, 2003; Ivessa *et al*, 2002), or stable DNA structures such as G-quadruplexes (Kumar *et al*, 2021; Paeschke *et al*, 2011; Ribeyre *et al*, 2009; Williams *et al*, 2023). Previous work showed that both Rrm3 and Pif1 are important for the viability of *dia2Δ* cells (Blake *et al*., 2006; Morohashi *et al*., 2009), and we found that *dia2Δ* cells lacking both Rrm3 and Pif1 are inviable (Fig 5A). Moreover, the *mcm7-10R rrm3Δ pif1Δ* triple mutant is also inviable (Fig 5B). In contrast, other replication fork associated helicases such as Chl1 (Samora *et al*, 2016; Skibbens, 2004; Srinivasan *et al*, 2020), Srs2 (Arbel *et al*, 2020; Crickard & Greene, 2019; Lehmann *et al*, 2020) and the Sgs1 orthologue of human BLM helicase (Gupta & Schmidt, 2020; Simmons *et al*, 2021) were not required for the viability of *mcm7-10R* (Fig EV4B-D). These findings indicate that Pif1-family helicases are important for cell viability in response to defects in CMG helicase disassembly.

**Figure 5.**
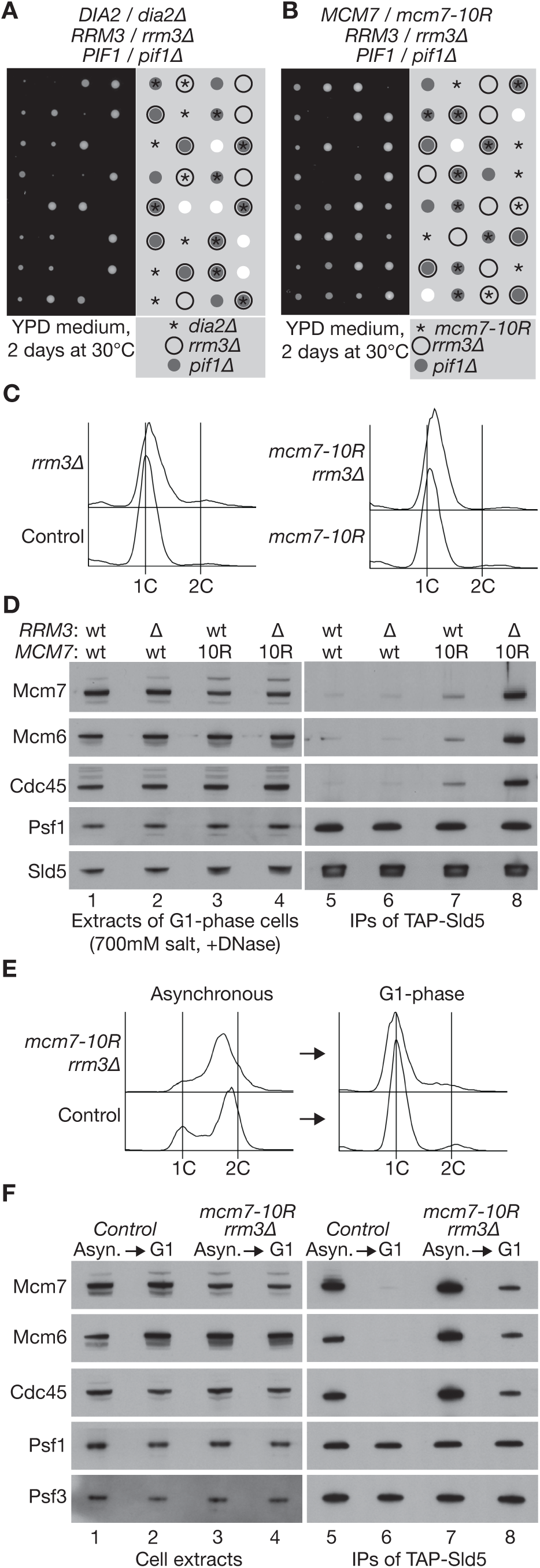
Pif1 family helicases are essential for viability and important for CMG helicase disassembly when Mcm7 ubiquitylation is defective. (**A**-**B**) Tetrad analysis for diploid yeast cells of the indicated genotypes. YPD = medium based on Yeast Extract, Peptone, Dextrose. (**C**) The indicated budding yeast strains (YSS47, YTM371, YCPR79 and YCPR143) were grown at 30°C and arrested in G1-phase by addition of mating pheromone. DNA content was monitored by flow cytometry. (**D**) Cell extracts were prepared from the samples in (C) and used to isolate the CMG helicase by immunoprecipitation of TAP-Sld5. The indicated factors were monitored by immunoblotting. (**E**) Control cells (YSS47 and *mcm7-10R rrm3Δ* (YCPR143) were grown asynchronously at 30°C and then arrested in G1-phase as above. (**F**) GINS and associated factors were isolated from cell extracts by immunoprecipitation of TAP-Sld5 and then monitored by immunoblotting.

To explore whether Rrm3 and Pif1 contribute to the disassembly of old CMG complexes, in cells in which the ubiquitylation of CMG-Mcm7 is defective, we initially tried to use degron technology to generate a conditional version of the lethal *mcm7-10R rrm3Δ pif1Δ* or *dia2Δ rrm3Δ pif1Δ* strains. However, this was not successful and so we focussed on characterising viable combinations of *mcm7-10R* with either *rrm3Δ* or *pif1Δ*, after arresting cells in G1-phase (Fig 5C). This revealed a striking persistence of the CMG helicase in the *mcm7-10R rrm3Δ* double mutant (Fig 5D, lane 8, compared to control cells (Fig 5D, lane 5), or to the *rrm3Δ* or *mcm7-10R* single mutants (Fig 5D, lanes 6-7).

Only a proportion of CMG helicase complexes persisted into G1-phase in the *mcm7-10R rrm3Δ* mutant (Fig 5E-F), either due to residual ubiquitylation of CMG-Mcm7-10R by SCF^Dia2^, or else reflecting the ability of additional factors to contribute to CMG helicase disassembly. Pif1 is a likely candidate, especially since it is essential for the viability of *mcm7-10R rrm3Δ* cells (Fig 5B). However, old CMG complexes did not persist during G1-phase in *mcm7-10R pif1Δ* cells (Fig EV4E-F), suggesting that the role of Pif1 is likely masked to some degree by the dominant role of Rrm3 in this pathway. In addition, CMG did not persist into G1-phase when *mcm7-10R* was combined with *chl1Δ*, *sgs1Δ* or *srs2Δ* (Fig EV4G-I). These findings highlight the dominant role of Rrm3 in the disassembly of old CMG helicase complexes that have not been removed via Mcm7 ubiquitylation and Cdc48.

### Rrm3 acts during S-phase to trigger disassembly of old CMG complexes from the previous cell cycle

To determine when Rrm3 acts during the cell cycle, to disassemble old CMG helicase complexes that have not been ubiquitylated by SCF^Dia2^, we generated *mcm7-10R rrm3Δ* cells that expressed the *RRM3* coding sequence at the *leu2* locus, under control of the regulatable *GAL1,10* promoter. Subsequently, *mcm7-10R rrm3Δ GAL-RRM3* cells were grown in the absence of Rrm3, in medium containing raffinose as the carbon source, before arresting in G1-phase by addition of mating pheromone. The culture was then split in two and one half switched to medium containing galactose and mating pheromone, to induce expression of *GAL-RRM3* for 60 minutes whilst maintaining arrest in G1-phase. Subsequently, the cultures were washed into fresh medium lacking mating pheromone, to allow cells to enter S-phase. Forty minutes later, mating pheromone was added once again, to arrest cells in G1-phase of the next cell cycle (Fig 6A). DNA content was monitored by flow cytometry throughout the experiment (Fig 6B), and samples corresponding to the first and second G1-phases were used to prepare cell extracts and monitor the presence of old CMG helicase complexes, by immunoprecipitation of TAP-tagged Sld5 (Fig 6C).

**Figure 6.**
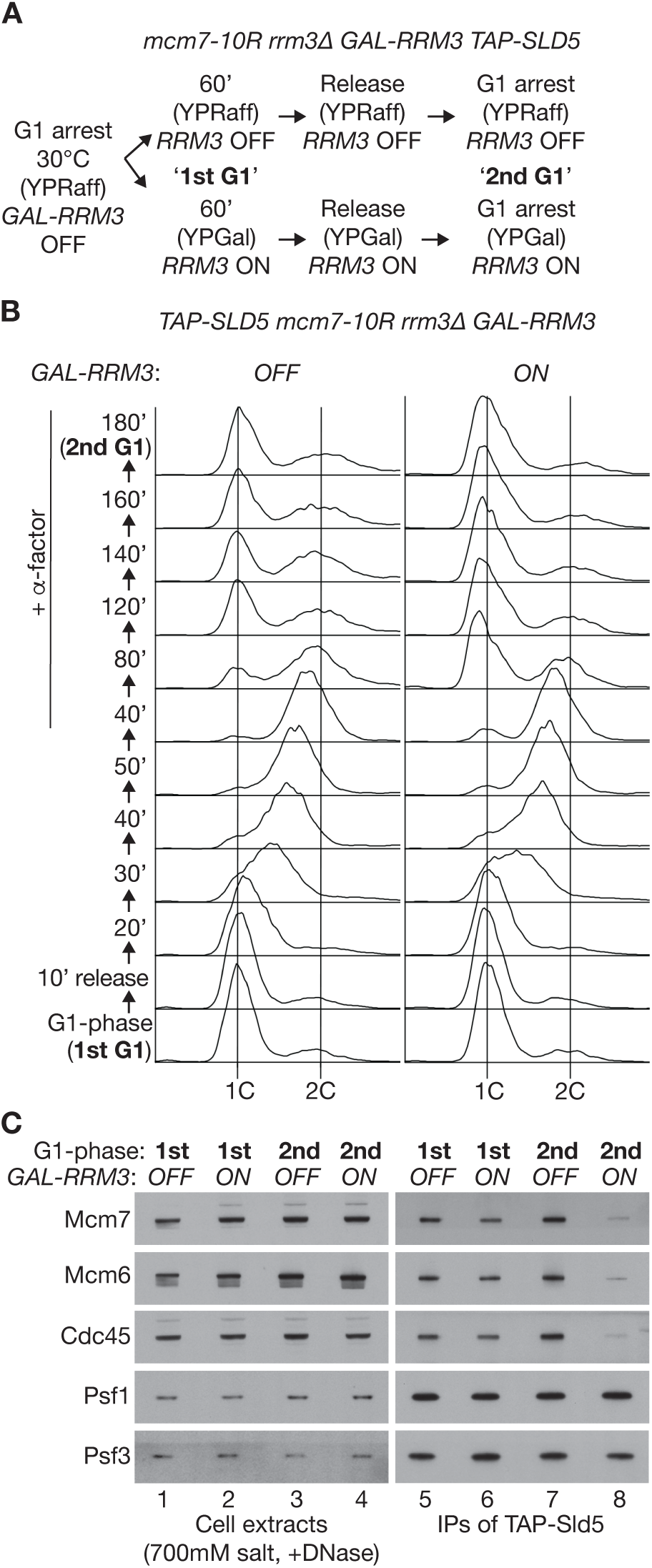
Rrm3 promotes disassembly of old CMG complexes after G1-phase. (**A**) Reaction scheme involving *mcm7-10R rrm3Δ GAL-RRM3 TAP-SLD5* cells (YCPR334. (**B**) DNA content from the experiment in (A) was monitored by flow cytometry. Upon release from the first arrest in G1-phase, mating pheromone (α-factor) was added again from 40 minutes onwards, to ensure that cells arrested subsequently in G1-phase of the second cell cycle. (**C**) Cells arrested in G1-phase arrest during the first and second cell cycles of the experiment in (B) were used to prepare cell extracts. The presence of CMG was monitored by immunoprecipitation of TAP-Sld5 and immunoblotting.

Induction of *GAL-RRM3* during G1-phase did not reduce the level of old CMG complexes in *mcm7-10R rrm3Δ GAL-RRM3* cells (Fig 6C, compare lanes 5-6). In contrast, when Rrm3 was present upon release from G1-phase and throughout the ensuing cell cycle, most old CMG complexes were disassembled (Fig 6C, compare lanes 7-8). This indicated that Rrm3 cannot drive the disassembly of old CMG complexes from G1 phase and instead acts at some later point in the cell cycle.

To test whether Rrm3 acts during S-phase to disassemble old CMG complexes from the previous cell cycle, the above experiment with *mcm7-10R rrm3Δ GAL-RRM3* cells was repeated, except that CMG was monitored at 10-minute intervals upon release from G1-phase (Fig 7A-D). In the absence of Rrm3 expression, the level of CMG increased 20 minutes after release from G1-phase and then persisted throughout S-phase, reflecting the addition of new CMG helicase complexes to the pool of old CMG from the previous cell cycle (Fig 7C, compare lanes 5-8), under conditions where CMG ubiquitylation was defective (Fig EV5A, *GAL-RRM3* **OFF** throughout experiment). The level of CMG also increased when *mcm7-10R rrm3Δ GAL-RRM3* cells entered S-phase in the presence of Rrm3 (Fig 7D, lanes 5-7). However, the amount of the CMG helicase peaked at 30 minutes and then declined 10 minutes later (Fig 7D, compare lanes 7-8). These data indicate that Rrm3 acts during S-phase to remove old CMG helicase complexes from the previous cell cycle. At the same time, new CMG complexes are assembled and then subsequently persist after DNA replication termination, due to the ubiquitylation defect of *mcm7-10R* cells (Fig EV5A, *GAL-RRM3* switched **ON** during G1-phase). In summary, these findings show that budding yeast cells have two pathways for CMG helicase disassembly, one of which works during DNA replication termination and involves ubiquitylation of Mcm7, whereas the second pathway acts during S-phase of the following cell cycle and is mediated by Pif1-family helicases and especially by Rrm3.

**Figure 7.**
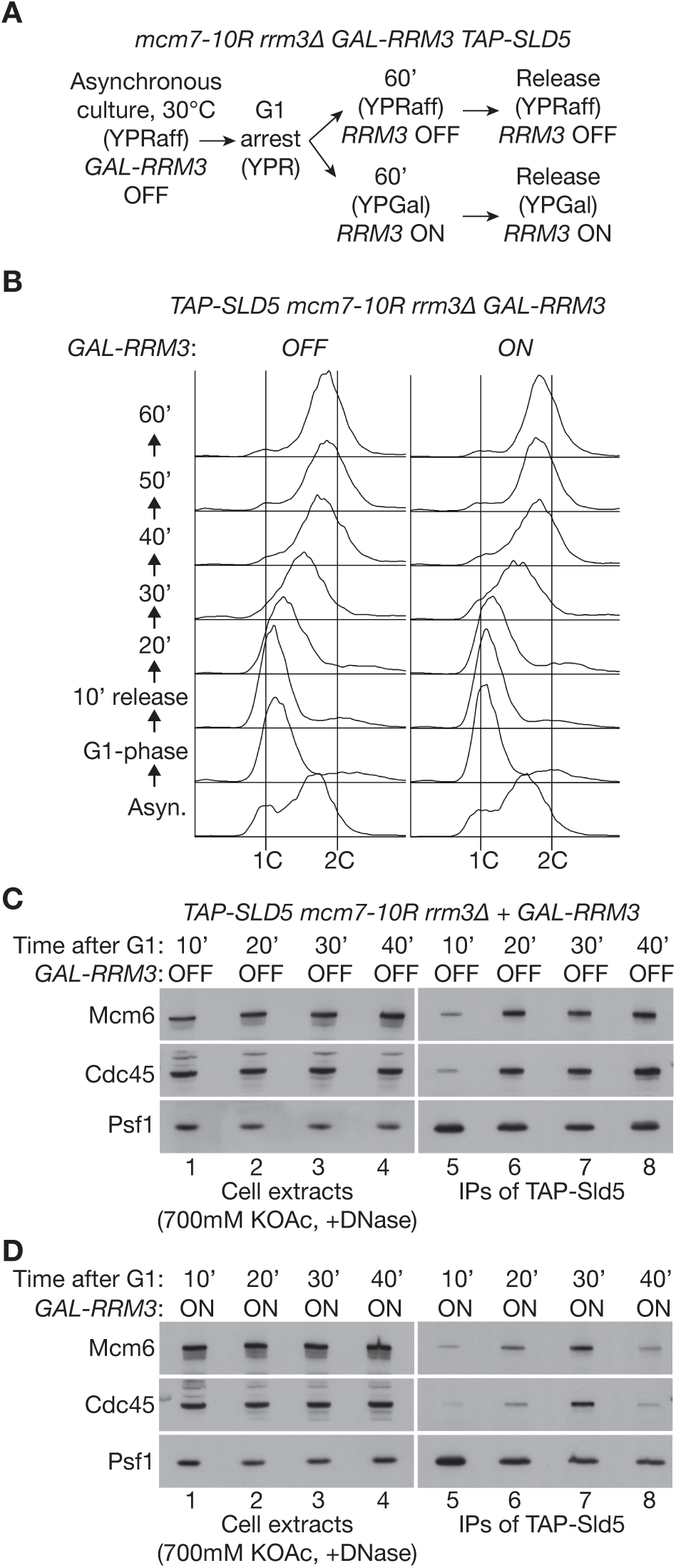
Rrm3 acts during S-phase to promote disassembly of old CMG complexes. (**A**) Reaction scheme involving *mcm7-10R rrm3Δ GAL-RRM3 TAP-SLD5* cells (YCPR334). (**B**) DNA content from the experiment in (A) was monitored by flow cytometry. (**C**) Samples of cells lacking Rrm3 (*GAL-RRM3* **OFF**) from the experiment in (B) were taken at the indicated times after release from G1-arrest and then used to prepare cell extracts. CMG was monitored by immunoprecipitation of TAP-Sld5 and immunoblotting. (**D**) Equivalent analysis for cells in which Rrm3 induction was induced after arresting cells in G1-phase (*GAL-RRM3* **ON**).

## Discussion

When two replication forks converge during DNA replication termination in budding yeast cells (Fig EV5B, steps (ii) to (iii)), the Rrm3 and Pif1 DNA helicases help to unwind the final stretch of parental DNA between the two replisomes (Claussin *et al*, 2022; Deegan *et al*., 2019), thereby facilitating the bypass of the two CMG helicases (Fig EV5B, step (iv)). Rrm3 is the major player in this process in budding yeast cells (Deegan *et al*., 2019), analogous to the dominant role of Rrm3 in the disassembly of old CMG complexes from the previous cell cycle (Figs 5-7; Fig EV4).

Rrm3 and Pif1 track along DNA in a 5’ to 3’ direction (Ivessa *et al*., 2002; Lahaye *et al*, 1993), and thus can associate with the opposite parental strand at replication forks to CMG. At a pair of converging replisomes, Rrm3-Pif1 migrate along the same parental strand as CMG from the opposing replisome (Fig EV5B), leading to a potential clash during DNA replication termination between Rrm3-Pif1 from one fork and CMG from the other. Nevertheless, Rrm3-Pif1 do not usually drive CMG disassembly under such conditions (Fig EV5B, step (iv)), and disintegration of the helicase is instead dependent upon SCF^Dia2^ and the Cdc48 unfoldase (Maric *et al*., 2014). In contrast, our data suggest that old CMG helicases from the previous cell cycle are disassembled by Rrm3-Pif1 during the subsequent S-phase (Fig EV5C, steps (ii) to (iii)), thereby allowing forks to converge and DNA synthesis to be terminated (Fig EV5B, steps (iv) to (v)).

The *mcm7-10R* allele delays CMG helicase disassembly and increases the proportion of cells with sub-nuclear foci of Rad52 in the following cell cycle (Fig 4G). This suggests that the encounter of new DNA replication forks with old CMG complexes increases the risk of DNA damage and is a major source of genome instability following defects in CMG helicase disassembly during DNA replication termination. The Rrm3-Pif1 pathway ameliorates the impact of old CMG helicase complexes on genome stability and is essential for cell viability when CMG ubiquitylation is defective in budding yeast cells. In the absence of both Rrm3 and Pif1, it is likely that old CMG complexes form a potent block to the progression or convergence of new DNA replication forks in the subsequent cell cycle. This is reminiscent of the encounter between a replication fork and an Mcm2-7 double hexamer followed by nucleosomes, which together form a barrier to fork progression that can be relieved by Pif1 helicase activity (Hill *et al*., 2020).

Our data indicate that the Rrm3-Pif1 pathway disrupts the CMG helicase into its component parts (Figs 5-7), rather than displacing intact helicase complexes from the DNA template. Defining the underlying mechanism is likely to require biochemical reconstitution of the encounter between Rrm3-Pif1 at a replication fork and old CMG. At this stage, multiple possibilities could be envisaged. For example, disassembly of old CMG complexes might be dependent on the force generated by two Rrm3-Pif1 helicase complexes, acting either side of an old CMG complex at a pair of converging replication forks. Alternatively, Rrm3-Pif1 might only trigger CMG disassembly when they encounter the C-terminal face of the Mcm2-7 ring of old CMG (Fig EV5C, step (ii), rightward forks). In this case, disassembly of old CMG might only require the encounter between a single replication fork and an old CMG complex. However, Rrm3-Pif1 would not be able to mediate CMG disassembly when forks converge in the absence of old CMG complexes, since Rrm3-Pif1 from one fork would encounter the amino-terminal face of Mcm2-7 from CMG at the second fork (Fig EV5B, step (iii). If Rrm3-Pif1 only disassemble CMG upon encountering the C-terminal face of Mcm2-7, then a similar mechanism might also explain how Pif1 helicases displace Mcm2-7 double hexamers during elongation (Hill *et al*., 2020), since this also involves an encounter between Pif1 and the C-terminal face of Mcm2-7.

A previous study of human CUL2^LRR1^ suggested that blocking CMG helicase disassembly during DNA replication termination in early replicons can interfere with CMG assembly at later origins, by preventing the recycling of CMG components such as CDC45 that might be present at limiting levels in some human cells (Fan *et al*., 2021). However, inactivation of CUL2^LRR1^ in *Xenopus* egg extracts does not affect the progression of DNA replication (Dewar *et al*., 2017; Sonneville *et al*., 2017), indicating that the sequestering on chromatin of limiting replisome components might only be an issue in specific cell types. In budding yeast, the efficiency of CMG assembly is similar in control cells and in cells lacking Dia2 (Maric *et al*., 2014), indicating that the sequestering of replisome components by old CMG complexes is unlikely to contribute to the genome instability that results from defects in helicase disassembly.

The existence of the Rrm3-Pif1 pathway has important implications for the evolution of CMG helicase disassembly in eukaryotic cells. The fact that fungi and animals ubiquitylate the Mcm7 subunit of CMG during DNA replication termination, using evolutionarily distinct ubiquitin ligases that are not conserved in plants, indicates that CMG ubiquitylation arose repeatedly during eukaryotic evolution. Therefore, ancestral eukaryotes must have disassembled the CMG helicase by alternative means that predated the ubiquitylation of Mcm7. Pif1-family helicases are broadly conserved in eukaryotic species and could have mediated such a pathway, acting during the subsequent S-phase to disassemble old CMG complexes from the previous cell cycle. The phenotypes of *mcm7-10R* indicate that delaying CMG to the subsequent cell cycle in ancestral eukaryotes would have come at the price of higher genome instability, thereby providing a selective pressure for the repeated emergence of CMG ubiquitylation, which then provided a mechanism for the rapid disassembly of CMG complexes during DNA replication termination in a single cell cycle.

Given the diversity of regulation between fungi and animal cells, it would be interesting in future studies to explore how CMG helicase disassembly is controlled in plants and in other eukaryotes for which chromosome duplication remains poorly characterised. It also remains to be determined whether Pif1-family helicases mediate CMG helicase disassembly in other species apart from budding yeast, and whether this role is shared with other 5’ to 3’ DNA helicases that help replication forks to bypass roadblocks, such as the RTEL1 helicase in metazoa (Hourvitz *et al*, 2023; Vannier *et al*, 2014).

## Methods

Reagents and resources from this study are listed in Appendix Table S3 and are available from MRC PPU Reagents and Services (https://mrcppureagents.dundee.ac.uk) or upon request.

### Yeast strains and growth

Yeast strains employed in this study are based on the W303 background and are listed in Appendix Table S3. Cells were grown in Yeast Extract Peptone (YP) medium, consisting of 1% (w/v) Yeast extract and 2% (w/v) Peptone. The medium was further supplemented with either 2% glucose (YPD medium), 2% raffinose (YPR medium), or 2% galactose (YPG medium). Unless specified otherwise, yeast cultures were grown at 30°C.

To synchronize *MAT***a** haploid cells in the G1-phase of the cell cycle, mid-log cell cultures were treated with mating pheromone (α-factor) at a concentration of 7.5 μg/mL. After one-hour, additional aliquots of 2.5 μg/mL α-factor (or 5 μg/mL in the case of *dia2Δ* cells) were added every 15 minutes until more than 90% of cells were unbudded or exhibited ’shmoos’. To release from the G1 arrest into S-phase, cells were washed and resuspended in fresh media lacking the pheromone and then left to grow.

To arrest cells in early S-phase, cells were released from G1 arrest and transferred into fresh media supplemented with 0.2M hydroxyurea until approximately 90% of the cells had initiated bud formation.

Degradation of *cdc48-aid* was induced by addition of 0.5mM of the auxin 3-indoleacetic acid to the cell culture.

### Meiotic progeny analysis of diploid cells

Diploid yeast cells were induced to enter meiosis by culturing them in Rich Sporulation Media (RSM) containing 0.25% (w/v) yeast extract, 1.5% (w/v) potassium acetate, 0.1% (w/v) glucose, 2% (w/v) agar and 770mg/L ‘supplement mixture’ comprising 100mg/L each of adenine and uracil, 50mg/L each of L-histidine, L-leucine, L-methionine, L-arginine, L-lysine and L-tryptophan, 20mg/L L-Tyrosine and 250mg/L of L-phenylalanine. Ascus walls were digested for 20 minutes with β-glucuronidase and spores were then separated using a dissection microscope (Singer Instruments). After growth on a YPD plate for two days at 30°C, the genotype of each colony was checked by replica plating onto selective media.

### Analysis of yeast cell growth by serial dilution on solid medium

A single colony from each yeast strain was suspended in 1 mL of 1X PBS at pH 7.4. Subsequently, cells were counted and adjusted to a final cell density of 0.33 x 10^7^ cells/mL, and three consecutive 10-fold dilutions were prepared and vortexed.

Finally, 15μL drops of each dilution were spotted onto plates (the location of each drop abeing determined by a grid placed below the plate), resulting in a total of 5 x 10^4^, 5 x 10^3^, 5 x 10^2^, and 50 cells. After incubation and colony formation, the plates were imaged using an Epson Expression 10000XL scanner every 24 hours for up to 5 days, depending on the conditions.

### Flow cytometry analysis

To monitor DNA content, a 1mL aliquot of cells (∼10^7^ cells) was fixed by resuspending them in 70% (v/v) ethanol. Following fixation, cells were processed as described previously (Labib *et al*, 1999) and analysed with a FACSCanto II flow cytometer (Becton Dickinson), and FlowJo software. Samples were gated manually after data processing to exclude cell fragments and other particles.

### Antibody coupling to magnetic beads

To initiate the antibody coupling process, 425µL of magnetic Dynabeads M-270 epoxy beads (Thermo Scientific, cat. No.14302D) were placed in a magnetic rack, and any residual DMF storing solution was carefully removed from the beads. Subsequently, the beads were thoroughly washed with 1mL of 1M sodium phosphate pH 7.4 with agitation on a rotating wheel for ten minutes. Following this, the supernatant was discarded and the beads were additionally washed twice with sodium phosphate.

For antibody coupling, 320µg of anti-Sld5 antibody was combined with 300µL of 3M ammonium sulphate and the appropriate amount of 1M sodium phosphate pH 7.4, to give a total volume of 900µL. The mixture was then incubated for two days at 4°C with constant agitation.

Beads were then placed in a magnetic rack and the supernatant was removed. The beads were then washed four times with 1X PBS before a 10 minute incubation in PBS/0.5% NP-40. Following this step, the beads were incubated with PBS/BSA twice for five minutes, and finally resuspended in 900µL of PBS/BSA solution supplemented with 0.02% sodium azide.

### Immunoprecipitation of protein complexes from yeast cell extracts

CMG complexes were isolated from yeast cell cultures as described previously (De Piccoli *et al*, 2012; Gambus *et al*., 2006; Maric *et al*., 2014). Briefly, yeast cell pellets from a 250mL culture, were centrifuged at 200 g for 3 minutes. The initial pellet was washed with 50mL of Tris-Acetate pH 9, followed by a second wash with 10mL of lysis buffer (100 mM Tris-Acetate pH 9, 700 mM potassium acetate, 10 mM magnesium acetate, 2 mM EDTA). Cells were then pelleted, weighed, and suspended in 3 volumes of lysis buffer supplemented with 2 mM sodium fluoride, 2 mM sodium β-glycerophosphate pentahydrate, 1 mM dithiothreitol (DTT), 1% Protease Inhibitor Cocktail (P8215, Sigma-Aldrich), and 1× Complete Protease Inhibitor Cocktail (05056489001, Roche).

The suspended cells were frozen dropwise in a 50mL Falcon tube filled with liquid nitrogen, resulting in the formation of ’yeast popcorn’. After allowing the liquid nitrogen to evaporate, the popcorn was stored at -80°C. Subsequently, equal amounts of popcorn for each sample were ground in a SPEX SamplePrep 6780 Freezer/Mill, using two cycles at ’rate 14’ (each cycle consisting of 2 minutes of ’run’ and 2 minutes of ’cool’). The resulting yeast cell powder was recovered and allowed to thaw at room temperature. Cell extracts were then transferred to centrifuge tubes on ice, with volume measurements taken along the way. Each extract was then diluted with a 1/4 volume of lysis buffer supplemented with 50% glycerol, 700 mM potassium acetate, 10 mM magnesium acetate, 0.5% IGEPAL CA-630, 2mM EDTA, inhibitors, and DTT at the concentrations mentioned above. Subsequently, 800 U/mL of DNase (Pierce Universal Nuclease) was added to each extract, followed by a 30-minute incubation at 4°C on a rotation wheel to release CMG complexes from chromatin. Insoluble cell debris was then removed by centrifugation at 25,000 g for 30 minutes, followed by ultracentrifugation at 100,000 g for 1 hour.

From each of the resulting extracts, a 50μL aliquot was mixed with 100μL 1.5X Laemmli Buffer and heated at 95°C for 2 minutes. The remainder of each extract was combined with 100μL antibody-coupled magnetic beads (approximately 1.7 x 10^9^ beads), ensuring that a fixed volume of extract was used for every sample in each experiment. Immunoprecipitation of protein complexes occurred over a 2-hour period at 4°C on a rotation wheel. Following incubation, proteins bound to magnetic beads were washed four times with 1mL of IP Wash Buffer (100 mM Tris-Acetate pH 9, 100 mM potassium acetate, 10 mM magnesium acetate, 2 mM EDTA, 0.1% IGEPAL CA-630). Eluted protein complexes were then separated from beads by adding 50μL of 1X Laemmli Buffer and heating the suspension at 95°C for 5 minutes. The supernatant was then removed and stored at -80°C until immunoblotting analysis.

### Mass spectrometry analysis

Samples were purified from yeast cell extracts synchronized in mid S-phase as above and eluted in 40 µl Laemmli buffer, of which 35 µl was resolved by SDS-polyacrylamide gel electrophoresis (SDS-PAGE) using NuPAGE Novex 4-12% Midi Bis-Tris gels (NP0321, Life Technologies) with NuPAGE MOPS SDS buffer (NP000102, Life Technologies). Subsequently, gels were stained with ‘SimplyBlue SafeStrain’ colloidal Coomassie (LC6060, Invitrogen) following the manufacturer’s instructions, and each lane was cut into 40 slices that were digested with trypsin before processing for mass spectrometry (MS Bioworks, USA). Data were analyzed using Scaffold software (Proteome Software Inc, USA). The raw mass spectrometry data for the experiments in Dataset EV1 and Dataset EV2 were submitted to the PRIDE database (https://www.ebi.ac.uk/pride/) with the project accession number PXD048935 and the project DOI 10.6019/PXD048935.

### Purification of recombinant CMG helicase complexes

Recombinant CMG helicase complexes were generated in budding yeast cells by co-expression of the 11 subunits from the bidirectional *GAL1,10* promoter. Each coding sequence was codon optimised to facilitate high-level expression, as described previously (Frigola *et al*, 2013), and the yeast Gal4 transcription factor was also co-expressed to enhance expression from the *GAL1,10* promoter. Cells with integrated plasmids expressing the 11 subunits of a particular version of CMG were cultured at 30°C in YPR media until reaching a density of 2-3 x 10^7^ cells/mL. Expression from the *GAL1,10* promoter was then induced by adding 2% galactose to the media, and cultures were allowed to grow at 30°C for 3 hours. Typically, 12 litres of cells were utilized for each CMG purification.

Cells were harvested by centrifugation at 1,935 g for 10 minutes, and the pellets were washed once with CMG Wash Buffer (25mM HEPES-KOH pH 7.6, 10% (v/v) glycerol, 300mM KCl, 2mM MgOAc, 0.02% (v/v) Tween-20, 1mM DTT), before resuspension in 0.3 volumes (relative to pellet mass) of CMG Buffer (same composition as CMG Wash Buffer but including 1X Complete Protease inhibitor cocktail). Cell suspensions were frozen dropwise in a 500mL container filled with liquid nitrogen, resulting ’yeast popcorn’, as described above. This material was ground to a fine powder using a SPEX CertiPrep 6850 FreezerMill for four cycles of 2 minutes, each at a rate of 15. Each cycle consisted of 2 minutes of ’run’ and 2 minutes of ’cool,’ and the resulting powder was stored at -80°C.

Subsequently, the frozen powder was thawed at room temperature and mixed with one volume of CMG Buffer. The mixture was then centrifuged at 235,000 g for 1 hour at 4°C. The soluble extract, typically around 40mL, was recovered and mixed with 4mL of anti-FLAG M2 affinity resin for 3 hours at 4°C on a rotating wheel. Subsequently, the resin was collected and extensively washed (typically in 20 column volumes) with CMG Buffer, and an additional column volume with CMG Wash Buffer. CMG complexes were then eluted by incubating the beads for 30 minutes on ice in 1 column volume of CMG FLAG Elution Buffer (same composition as CMG Wash Buffer supplemented with 0.5 mg/mL FLAG peptide), and the eluted material then separated from the beads. This step was repeated by incubating the beads with CMG FLAG Elution Buffer that contained 0.25mg/mL FLAG peptide, and both eluted samples were pooled in the same tube.

FLAG eluates were then adjusted to 2mM CaCl_2_ and mixed with 3mL of Calmodulin affinity resin before incubation for 1 hour at 4°C on a rotating wheel. The beads were then collected and mixed with 10mL of CMG Calmodulin Elution Buffer (25mM HEPES-KOH pH 7.6 10% (v/v) glycerol, 300mM KCl, 2mM MgOAc, 0.02% (v/v), Tween-20, 2mM EDTA, 2mM EGTA and 1mM DTT). Subsequently, the eluates were collected and pooled.

The eluate fraction was loaded onto a 0.2mL MiniQ column (GE Healthcare Lifesciences) in CMG Wash Buffer. CMG complexes were eluted from the MiniQ column using a 4mL gradient ranging from 0.3M to 0.6M KCl, which was automatically mixed by an Akta system. After elution, fractions were analysed by SDS-PAGE followed by Coomassie staining, and those containing CMG were pooled and dialyzed overnight at 4°C against CMG Dialysis Buffer using a Slide-A-Lyzer dialysis cassette (Thermo). The dialyzed sample was recovered, aliquoted, and snap-frozen. The final concentration of CMG complexes was determined by SDS-PAGE followed by Coomassie staining, comparing it with a titration of known concentrations of Bovine Serum Albumin (BSA).

### *In vitro* CMG ubiquitylation assay

To assemble reconstituted reactions for the CMG ubiquitylation process, ubiquitylation enzymes were purified using previously established protocols (Deegan et al., 2019; Deegan et al., 2020; Yeeles et al., 2015).

The reactions, typically 8μL in volume, comprised 15nM CMG, 30nM Uba1 (E1 enzyme, provided by Axel Knebel, MRC PPU), 15nM Cdc34 (E2 enzyme), 1nM or 25nM SCF^Dia2^ (E3 enzyme), 30nM Ctf4, 30nM Pol ε, 45nM Mrc1, 6μM Ubiquitin (depending on the experiment, either wild-type ubiquitin, or lysine-free ubiquitin known as K0-Ubiquitin) and 5mM ATP. Reactions were assembled on ice in Reaction Buffer (25mM HEPES-KOH pH 7.6, 75mM KOAc, 10mM MgOAc, 0.02% IGEPAL CA-630, 0.1mg/mL BSA, 1mM DTT) and then incubated for 20 minutes at 30°C, unless stated otherwise. To halt the reactions, 2X LDS Buffer (ThermoFisher Scientific) was added, and the mixture was boiled at 95°C for 5 minutes.

For the experiment in Figure EV2D, ubiquitylation reactions were performed as mentioned above and each sample was incubated at 4°C with 2.5µL of magnetic beads coupled to anti-Cdc45 antibody. Protein mixtures were incubated for 1 hour and CMG complexes bound to the beads were washed twice with 190µL of Reaction Buffer containing 150mM KOAc. Finally, beads were resuspended in 1X LDS and boiled for 5 minutes at 95°C.

In experiments involving Mcm7 with inserted TEV cleavage sites, CMG ubiquitylation reactions were performed in Reaction Buffer containing NaOAc instead of KOAc, and then stopped by adding 700mM NaOAc to the reaction mix. Subsequently, the mixture was incubated with 0.5 mg/mL TEV protease (provided by Axel Knebel, MRC PPU) for 40 minutes at 30°C.

### *In vitro* CMG disassembly assay

The Cdc48 segregase and the Ufd1-Npl4 heterodimer were purified as described previously (Deegan *et al*., 2020). Following ubiquitylation reactions as above, 50nM Cdc48 and 50nM Ufd1-Npl4 were added and incubation continued for an additional 20 minutes at 30°C. Subsequently, 700mM NaOAc was added to stop the reaction before addition of 2.5μL of magnetic beads coupled with an antibody recognizing the Sld5 subunit of the CMG helicase. The mixture was incubated for 40 minutes at 4°C with constant shaking at 1,400 rpm on an Eppendorf Thermomixer. Subsequently, bound proteins were washed twice with Reaction Wash buffer (25mM HEPES-KOH pH 7.6 150mM KOAc, 10mM MgOAc, 0.02% (v/v) IGEPAL CA-630, 0.1mg/mL BSA and 1mM DTT), and then eluted by addition of 2X LDS Buffer and heating at 95°C for 5 minutes.

### *In vitro* assay to monitor SCF^Dia2^ interaction with CMG

To assess the association of SCF^Dia2^ with CMG complexes, 20μL reactions were prepared as for the ubiquitylation reactions described above, except that the SCF^Dia2^ concentration was increased to 10nM to facilitate detection by immunoblotting, and ubiquitin and ATP were excluded. Reactions were incubated for 15 minutes on ice and input samples (5μL) then collected. Subsequently, 3.75μL of magnetic beads coupled to anti-Sld5 antibodies were added to the remainder of each reaction, before a 30-minute incubation at 4°C with constant shaking at 1,400 rpm on an Eppendorf Thermomixer. Bound CMG complexes were recovered, washed twice in Reaction Wash buffer and then eluted by adding 2X LDS Buffer and heating at 95°C for 5 minutes.

### SDS-PAGE and immunoblotting

Proteins were separated on NuPAGE Novex 4-12% Bis-Tris gels with NuPAGE MOPS SDS buffer. For better resolution of ubiquitin chains, NuPAGE 3– 8% Tris-Acetate gels were used in combination with NuPAGE Tris-Acetate SDS buffer. Subsequently, proteins were either stained with colloidal Coomassie blue dye or transferred to nitrocellulose membrane utilizing the iBlot Dry Transfer System (Life Technologies) in accordance with the manufacturer’s instructions.

The antibodies used to detect the proteins in this study are listed in Appendix Table S3.

### Detection of sub-nuclear foci of Rad52-GFP foci in yeast cells

To detect Rad52-GFP foci, 1mL of yeast cell cultures were harvested and fixed by mixing with 1 mL 16% paraformaldehyde solution (Thermo Scientific, cat. No. 043368), to give a final concentration of 8% Paraformaldehyde. The mixture was incubated for 10 minutes at room temperature on a rotating wheel and subsequently washed in 1X PBS. After the wash, cells were resuspended in 1mL 1X PBS and stored at 4°C. To visualize DNA, cells were pelleted at 845 g for 3 minutes and incubated for 30 minutes at room temperature in 500μL of 1μg/mL DAPI. Subsequently, cells were washed three times in 1X PBS to reduce background fluorescence and then finally resuspended in a suitable volume for microscopy (typically 20μL). A 4μL aliquot was then placed onto a microscope slide and mounted with a coverslip sealed with nail polish. Cells were analysed using a Yokogawa CSU-X1 spinning disk microscope with a HAMAMATSU C13440 camera, equipped with a PECON incubator and a 63X/1.40 Plan Apochromat oil immersion lens (Olympus). Multiple fields were imaged for each strain. Images were captured using ZEN blue software (Zeiss), processed, and analyzed with ImageJ software (National Institute of Health).

Approximately 100 cells per strain were counted in three independent experiments, to determine the percentage with Rad52-GFP foci. Mean values were then calculated, together with the associated standard deviation. The samples were compared using a one-way ANOVA test followed by Tukey’s test correction to assess statistical significance, utilizing Prism9 software (GraphPad).

### Sequence alignment and structural analysis

For Fig EV2A, protein sequences were aligned with Clustal Omega software (https://www.ebi.ac.uk/Tools/msa/clustalo/) and the alignment displayed with MView (https://www.ebi.ac.uk/Tools/msa/mview/).

AlphaFold2-multimer (Mirdita *et al*, 2022) was used to generate the model in Fig S1B and Movie EV2 of SCF^Dia2^. Structural models were displayed, and Movies generated, using UCSF Chimera (Pettersen *et al*, 2021).

## Acknowledgements

We thank Ryo Fujisawa for help with protein purification and MRC PPU Reagents and Services (https://mrcppureagents.dundee.ac.uk) for antibody production. We are grateful for the support of the Medical Research Council (core grants MC_UU_12016/13 to KL and MC_UU_0035/4 to TD) and Cancer Research UK (Programme Grant C578/A24558 and PhD studentship C578/A25669 to KL). Materials generated in this study are listed in Appendix Table S3 and are available from MRC PPU Reagents and Services (https://mrcppureagents.dundee.ac.uk) or upon request.

## Disclosure and competing interests statement

The authors have no competing interests.

---

**Figure EV1.**
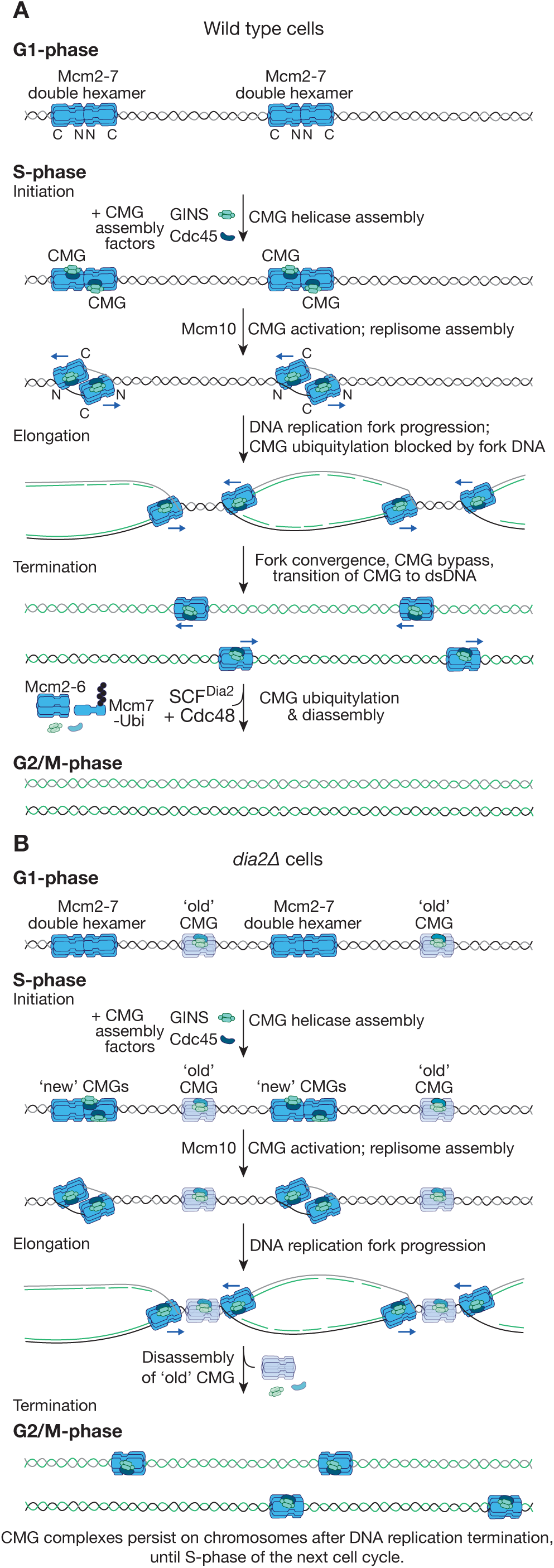
Model for disassembly of CMG helicase in wild type cells and dia2Δ. (**A**) In wild type budding yeast cells, ubiquitylation of CMG by SCF^Dia2^ is sterically impeded at replication forks, by the parental DNA strand that is excluded from the Mcm2-7 ring of the helicase. This inhibition is released during DNA replication termination, when a pair of forks converge and the two CMG helicases bypass each other, thereby breaking the association between CMG and the excluded parental DNA strand. (**B**) In *dia2Δ* cells, CMG cannot be ubiquitylated during DNA replication termination and persists on chromatin until the next cell cycle. However, once dia2Δ enter S-phase, the old CMG complexes are disassembled by a previously unknown pathway, likely coupled to the encounter between new replication forks and old CMG.

**Figure EV2.**
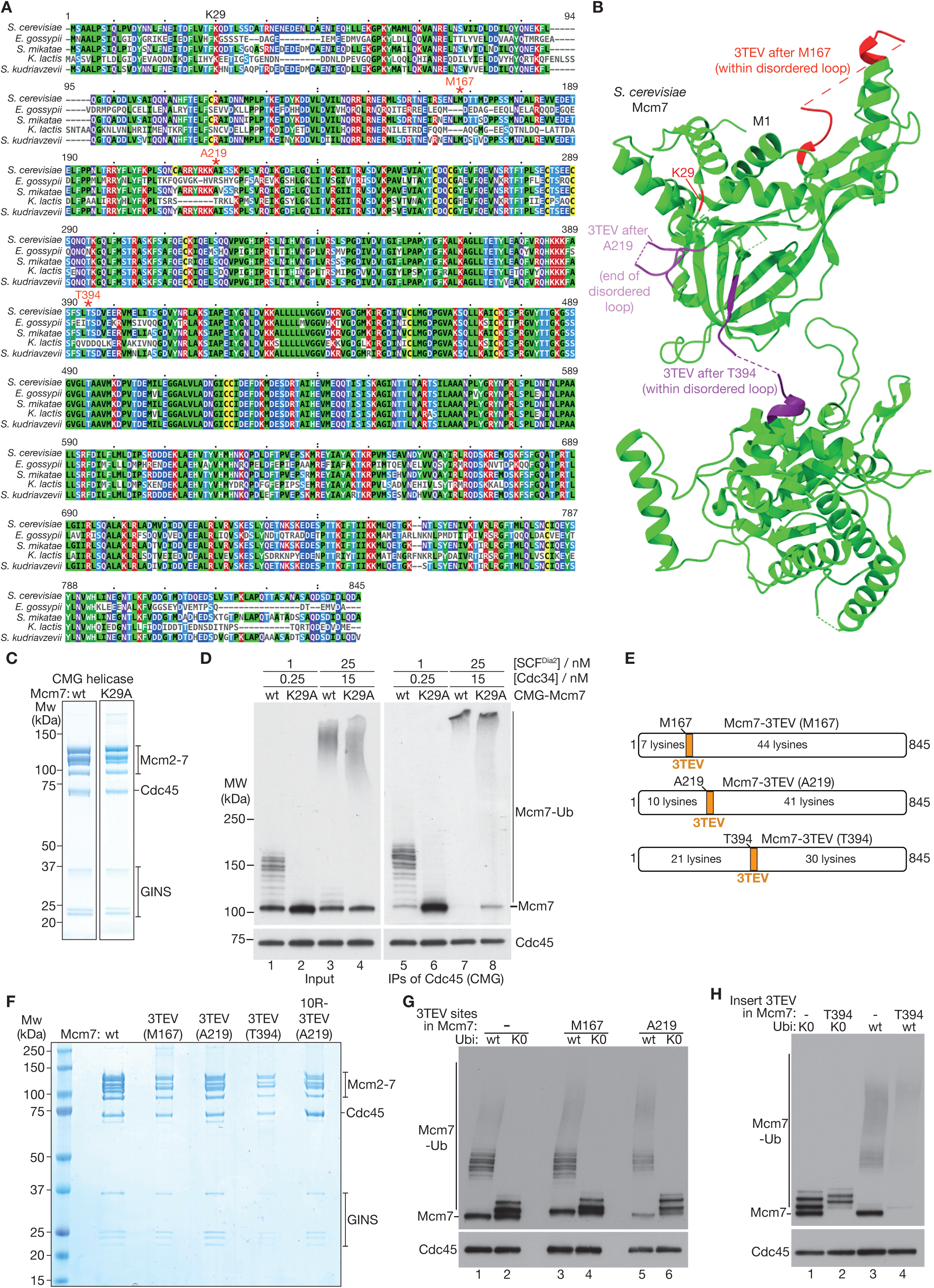
Generation and characterisation of recombinant CMG complexes with mutated alleles of Mcm7. (**A**) Mcm7 from multiple yeast species (*S. cerevisiae* = *Saccharomyces cerevisiae*; *E. gossypii* = *Eremothecium gossypii*; *S. mikatae* = *Saccharomyces mikatae*; *K. lactis* = *Kluyveromyces lactis*; *S. kudriavzevii* = *Saccharomyces kudriavzevii*) were aligned using Clustal Omega software. Mcm7-K29 is marked in black, whereas the three sites used for insertion of TEV cleavage sites (M167, M219 and M394) are shown in red with an asterisk. (**B**) The structure of *S. cerevisiae* Mcm7 (from PDB file 7PMK), illustrating the location of K29 and the three sites within disordered loops that were used to insert three consecutive TEV cleavage sites (after M167, A219 or T394 of Mcm7). Also see Movie EV1. (**C**) Purified recombinant CMG helicase, with wild type Mcm7 or Mcm7-K29A, were resolved by SDS-PAGE, before staining of the gel with Coomassie-blue. (**D**) CMG containing wild type (wt) Mcm7 or Mcm7-K29A was ubiquitylated in the presence of the indicated concentrations of SCF^Dia2^ and Cdc34. The reactions were analysed by immunoblotting. (**E**) Distribution of lysines either side of TEV cleavage sites inserted after M167, A219 or T394 of Mcm7 (3TEV = three consecutive TEV cleavage sites). (**F**) Coomassie-stained gel with purified recombinant CMG containing wild type Mcm7 or the indicated variants. (**G**-**H**) Recombinant CMG with the indicated versions of Mcm7 were ubiquitylated in vitro by SCFDia2 and Cdc34, using either wild type ubiquitin (wt Ubi) or lysine-free ubiquitin (K0 Ubi). Reactions were then analysed by immunoblotting.

**Figure EV3.**
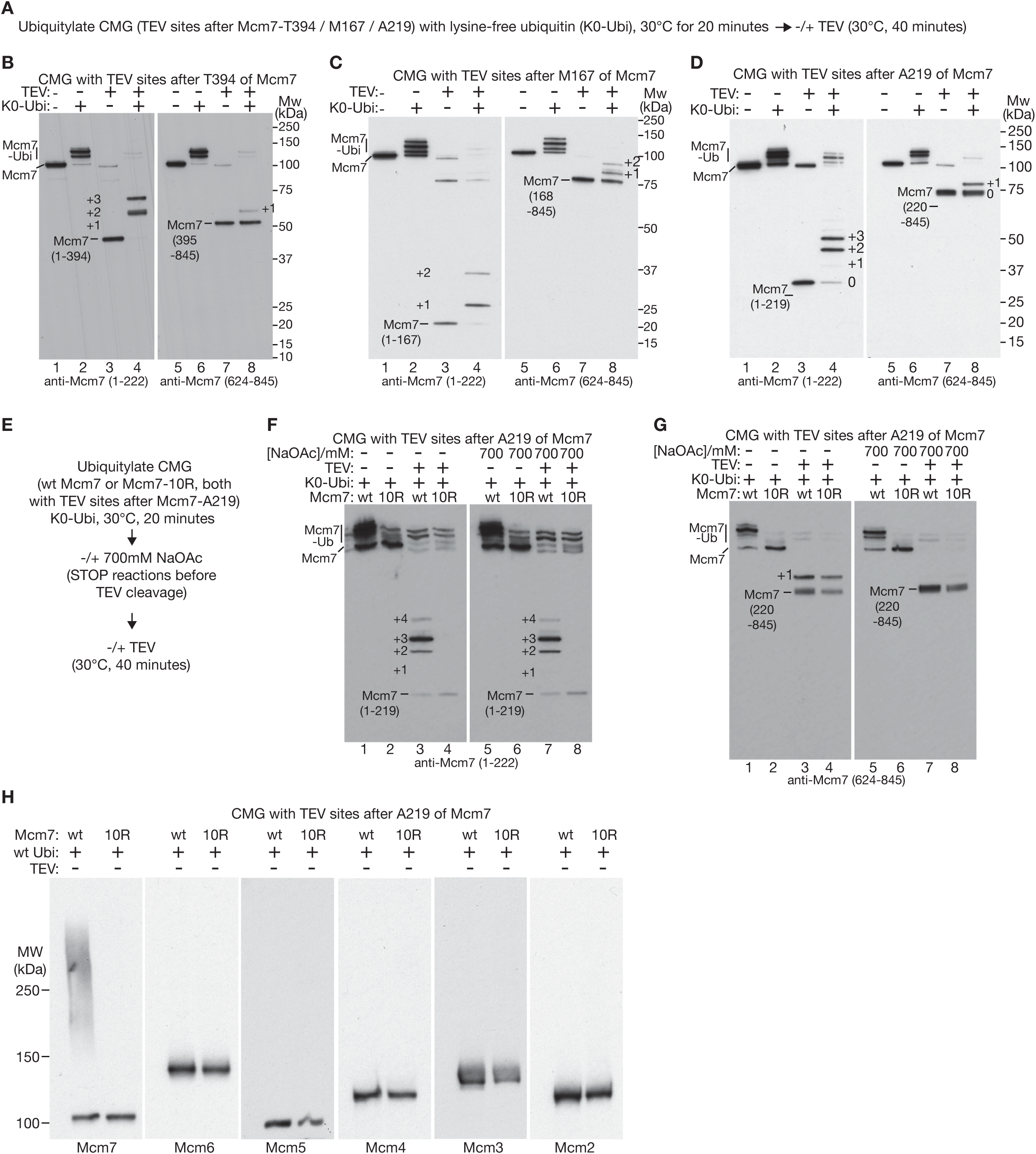
CMG ubiquitylation by SCF^Dia2^ and Cdc34 has a strong preference for the amino-terminal region of Mcm7. (**A**) Reaction scheme. (**B**) CMG with three TEV sites inserted after Mcm7-T394 was ubiquitylated in reactions containing lysine-free ubiquitin (K0-Ubi) as indicated. The reaction products were then cleaved with TEV protease as shown and analysed by immunoblotting with antibodies specific to the amino terminus of Mcm7 (1-222) or the carboxyl-terminus (624-845). (**C**-**D**) Equivalent analysis for CMG with three TEV sites inserted after Mcm7-M167 (C) or Mcm7-A219 (D). (**E**) Scheme for reconstituted CMG ubiquitylation reactions (K0-Ubi = lysine-free ubiquitin) that were subsequently stopped as indicated by increasing the salt concentration, before cleavage of TEV sites inserted after A219 of Mcm7 (see Methods for details). (**F**) Reactions described in (E) were analysed by immunoblotting with antibodies to Mcm7 1-222. (**G**) The same reactions were also analysed by immunoblotting with antibodies to Mcm7 624-845. (**H**) Analogous reactions to those in E-G were performed with wild type ubiquitin. The Mcm2-7 subunits of CMG were monitored by immunoblotting.

**Figure EV4.**
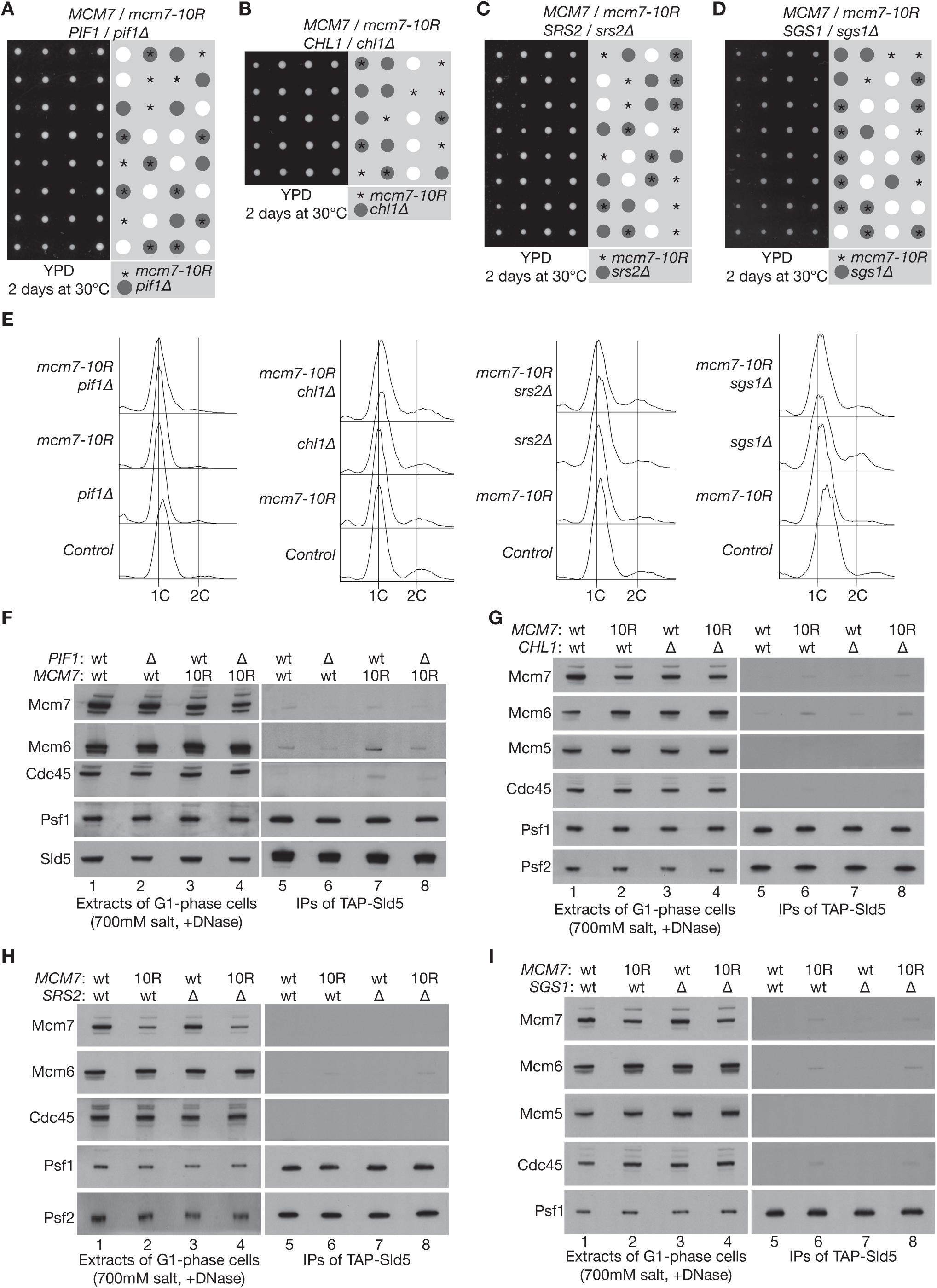
Combination of *mcm7-10R* with *pif1Δ*, *chl1Δ*, *srs2Δ* or *sgs1Δ* does not cause synthetic lethality or accumulation of CMG during G1-phase. (**A**-**D**) Tetrad analysis of diploid yeast cells of the indicated genotypes (YCPR75, YCPR435, YCPR434, YCPR171). YPD = Yeast Extract, Peptone, Dextrose. (**E**) The indicated strains (YCPR141, YCPR406, YCPR412, YCPR428) were arrested in G1-phase at 30°C by addition of mating pheromone. DNA content was monitored by flow cytometry. (**F**-**I**) Cell extracts were prepared from the samples in (E) and used to isolate GINS and the CMG helicase by immunoprecipitation of TAP-Sld5. The indicated factors were monitored by immunoblotting.

**Figure EV5.**
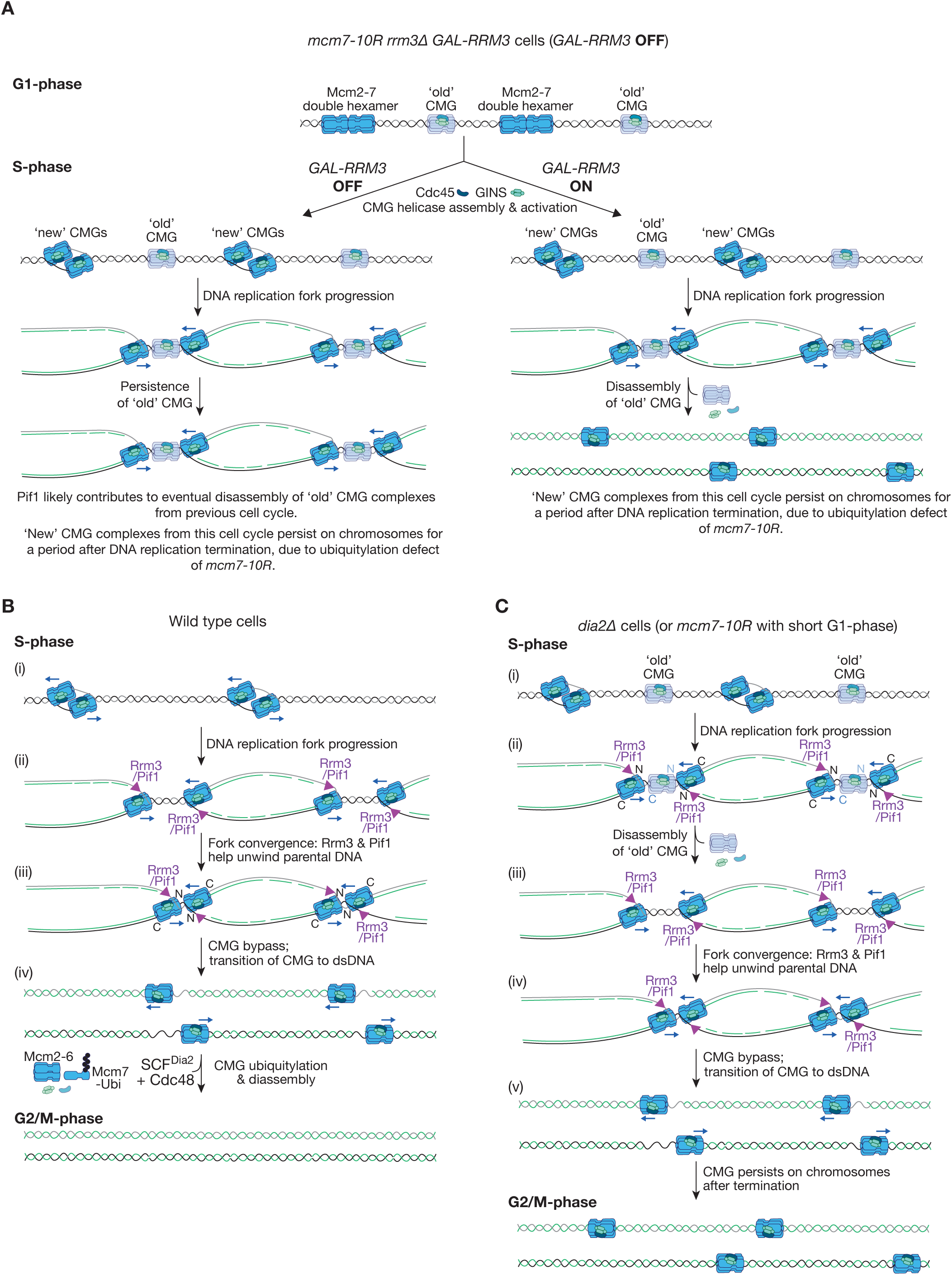
Models for disassembly of old CMG complexes by Rrm3-Pif1. (**A**) Scheme illustrating the behaviour of old and new CMG complexes for the experiment in Figure 7. (**B**) During DNA replication termination in wild type yeast cells, Rrm3 and Pif1 help to unwind the last stretch of parental DNA between converging forks during DNA replication termination, but do not trigger CMG helicase disassembly. See text for details. (**C**) *dia2Δ* cells (or *mcm7-10R* cells with a short G1-phase) enter S-phase with old CMG complexes still present on chromatin. These old CMG complexes are disassembled during S-phase, dependent upon Rrm3 (likely with backup from Pif1 – see text for further details). Subsequently, Rrm3-Pif1 help to unwind the last stretch of parental DNA between converging forks during DNA replication termination, leaving the new CMG complexes from the current cell cycle on chromatin, due to the defect in CMG ubiquitylation.

## Appendix – Table of Contents

Appendix Figure S1 Page 1

Appendix Figure S2 Page 2

Appendix Table S1 Page 3

Appendix Table S2 Page 5

Appendix Table S3 Page 7

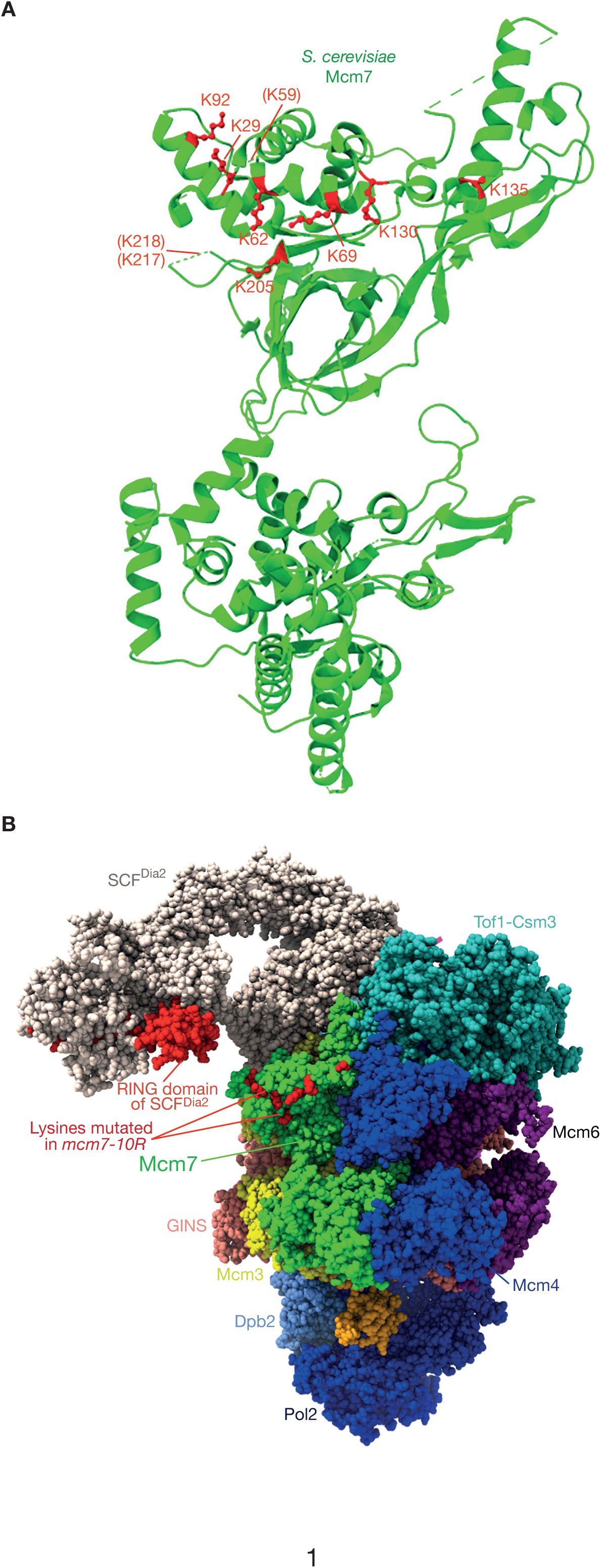
Polo Rivera et al_Appendix Figure S1

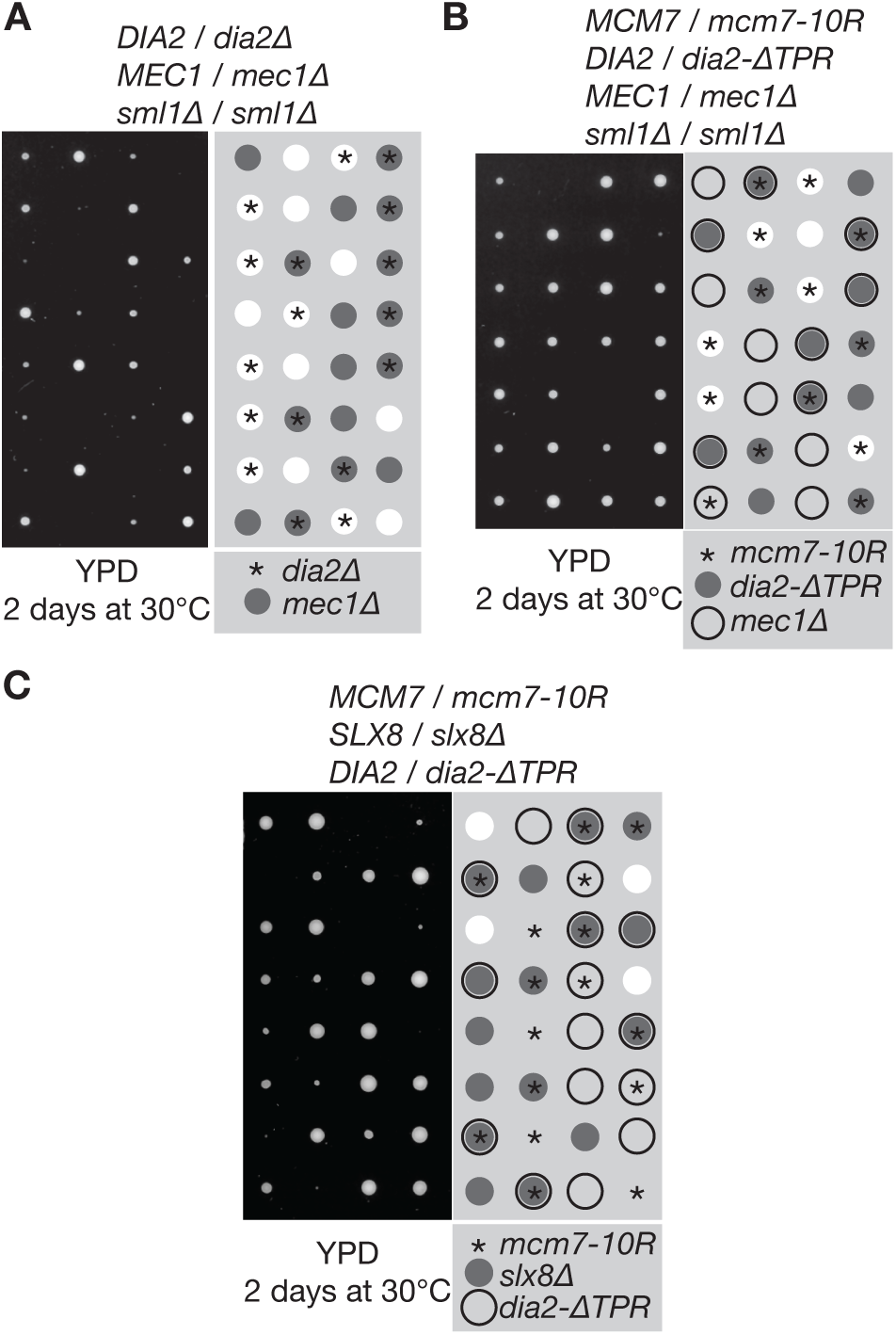
Polo Rivera et al_Appendix Figure S2

**Appendix Table S1.**
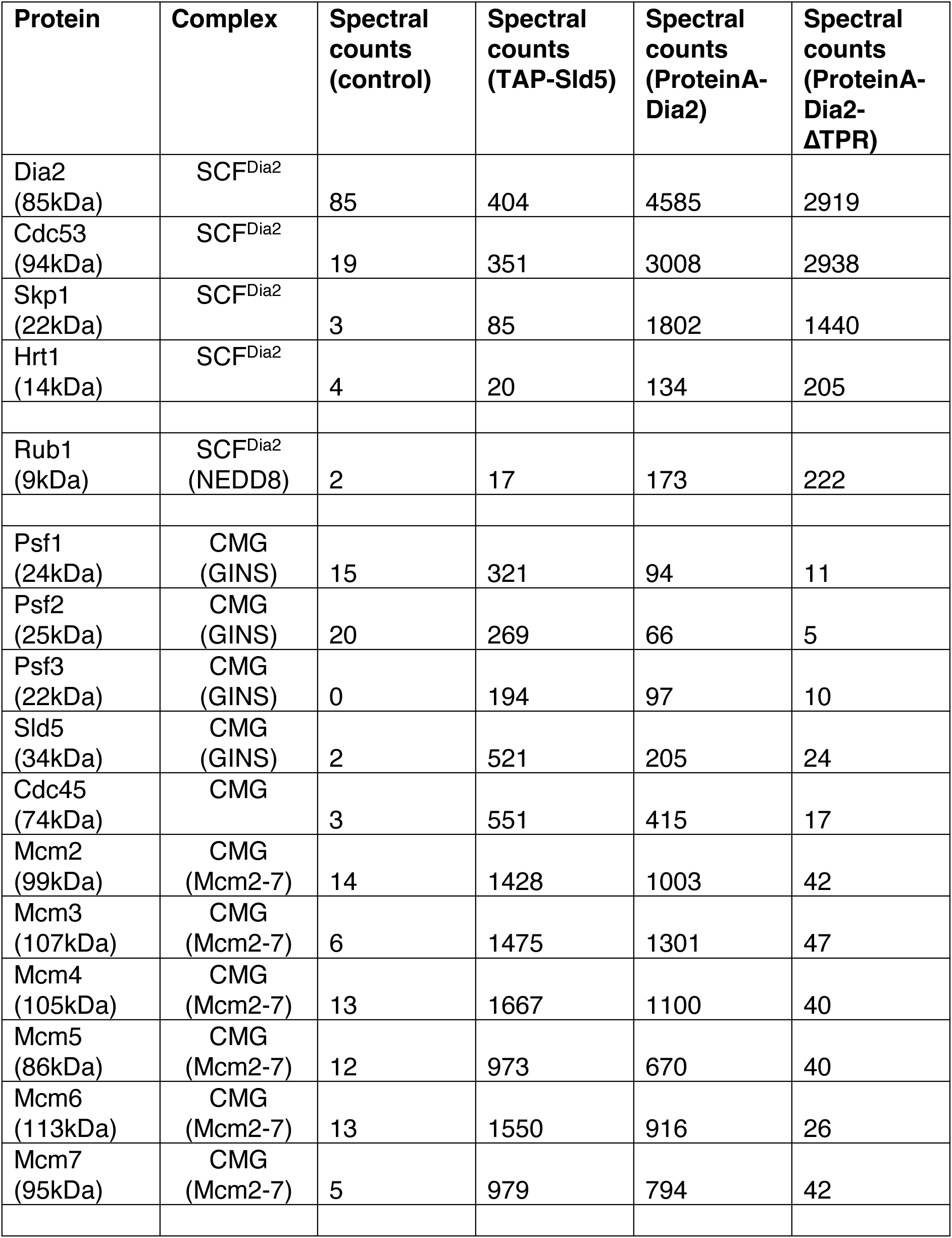
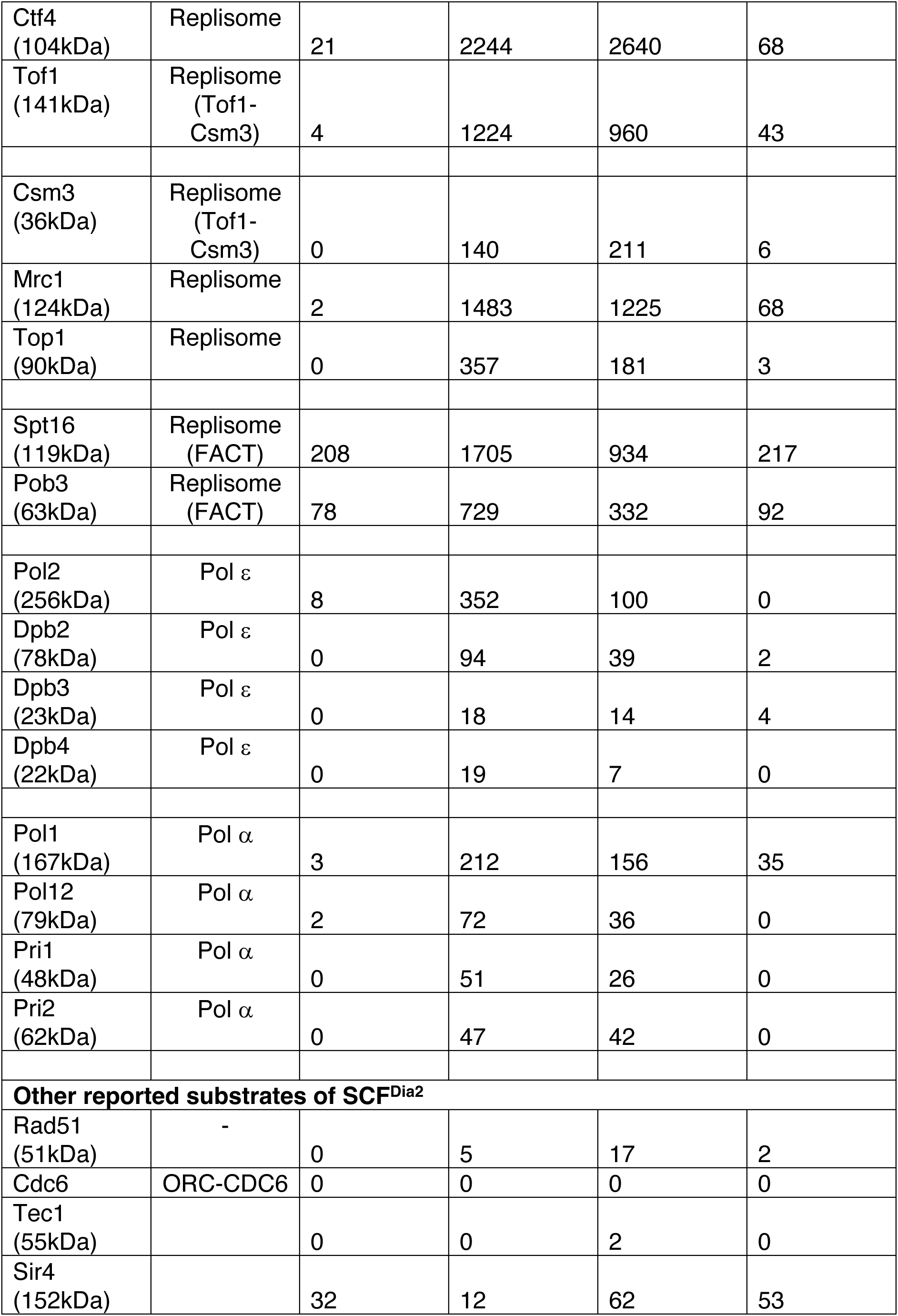
Mass spectrometry data showing that CMG and associated replisome components are the major partners of SCF^Dia2^, largely dependent on the TPR domain of Dia2. Also see Dataset EV1. This is the first of two independent experiments (the other is in Appendix Table S2).

| **Protein** | **Complex** | **Spectral counts (control)** | **Spectral counts (TAP-Sld5)** | **Spectral counts (ProteinA-Dia2)** | **Spectral counts (ProteinA-Dia2-ΔTPR)** |
| --- | --- | --- | --- | --- | --- |
| Dia2 (85kDa) | SCF<sup>Dia2</sup> | 85 | 404 | 4585 | 2919 |
| Cdc53 (94kDa) | SCF<sup>Dia2</sup> | 19 | 351 | 3008 | 2938 |
| Skp1 (22kDa) | SCF<sup>Dia2</sup> | 3 | 85 | 1802 | 1440 |
| Hrt1 (14kDa) | SCF<sup>Dia2</sup> | 4 | 20 | 134 | 205 |
| Rub1 (9kDa) | SCF<sup>Dia2</sup> (NEDD8) | 2 | 17 | 173 | 222 |
| Psf1 (24kDa) | CMG (GINS) | 15 | 321 | 94 | 11 |
| Psf2 (25kDa) | CMG (GINS) | 20 | 269 | 66 | 5 |
| Psf3 (22kDa) | CMG (GINS) | 0 | 194 | 97 | 10 |
| Sld5 (34kDa) | CMG (GINS) | 2 | 521 | 205 | 24 |
| Cdc45 (74kDa) | CMG | 3 | 551 | 415 | 17 |
| Mcm2 (99kDa) | CMG (Mcm2-7) | 14 | 1428 | 1003 | 42 |
| Mcm3 (107kDa) | CMG (Mcm2-7) | 6 | 1475 | 1301 | 47 |
| Mcm4 (105kDa) | CMG (Mcm2-7) | 13 | 1667 | 1100 | 40 |
| Mcm5 (86kDa) | CMG (Mcm2-7) | 12 | 973 | 670 | 40 |
| Mcm6 (113kDa) | CMG (Mcm2-7) | 13 | 1550 | 916 | 26 |
| Mcm7 (95kDa) | CMG (Mcm2-7) | 5 | 979 | 794 | 42 |
| Ctf4<br>(104kDa) | Replisome | 21 | 2244 | 2640 | 68 |
| Tof1<br>(141kDa) | Replisome<br>(Tof1-Csm3) | 4 | 1224 | 960 | 43 |
| Csm3<br>(36kDa) | Replisome<br>(Tof1-Csm3) | 0 | 140 | 211 | 6 |
| Mrc1<br>(124kDa) | Replisome | 2 | 1483 | 1225 | 68 |
| Top1<br>(90kDa) | Replisome | 0 | 357 | 181 | 3 |
| Spt16<br>(119kDa) | Replisome<br>(FACT) | 208 | 1705 | 934 | 217 |
| Pob3<br>(63kDa) | Replisome<br>(FACT) | 78 | 729 | 332 | 92 |
| Pol2<br>(256kDa) | Pol $\epsilon$ | 8 | 352 | 100 | 0 |
| Dpb2<br>(78kDa) | Pol $\epsilon$ | 0 | 94 | 39 | 2 |
| Dpb3<br>(23kDa) | Pol $\epsilon$ | 0 | 18 | 14 | 4 |
| Dpb4<br>(22kDa) | Pol $\epsilon$ | 0 | 19 | 7 | 0 |
| Pol1<br>(167kDa) | Pol $\alpha$ | 3 | 212 | 156 | 35 |
| Pol12<br>(79kDa) | Pol $\alpha$ | 2 | 72 | 36 | 0 |
| Pri1<br>(48kDa) | Pol $\alpha$ | 0 | 51 | 26 | 0 |
| Pri2<br>(62kDa) | Pol $\alpha$ | 0 | 47 | 42 | 0 |
| **Other reported substrates of SCF<sup>Dia2</sup>** |  |  |  |  |  |
| Rad51<br>(51kDa) | - | 0 | 5 | 17 | 2 |
| Cdc6 | ORC-CDC6 | 0 | 0 | 0 | 0 |
| Tec1<br>(55kDa) |  | 0 | 0 | 2 | 0 |
| Sir4<br>(152kDa) |  | 32 | 12 | 62 | 53 |

**Appendix Table S2.** Mass spectrometry data showing that CMG and associated replisome components are the major partners of SCF^Dia2^, largely dependent on the TPR domain of Dia2. Also see Dataset EV2. This is the second of two independent experiments (the other is in Appendix Table S1).

| **Protein** | **Complex** | **Spectral counts (control)** | **Spectral counts (TAP-Sld5)** | **Spectral counts (ProteinA-Dia2)** | **Spectral counts (ProteinA-Dia2-ΔTPR)** |
| --- | --- | --- | --- | --- | --- |
| Dia2 (85kDa) | SCF<sup>Dia2</sup> | 73 | 515 | 4917 | 3775 |
| Cdc53 (94kDa) | SCF<sup>Dia2</sup> | 47 | 372 | 2125 | 2078 |
| Skp1 (22kDa) | SCF<sup>Dia2</sup> | 5 | 47 | 1002 | 998 |
| Hrt1 (14kDa) | SCF<sup>Dia2</sup> | 4 | 14 | 160 | 132 |
| Rub1 (9kDa) | SCF<sup>Dia2</sup> (NEDD8) | 2 | 24 | 105 | 77 |
| Psf1 (24kDa) | CMG (GINS) | 0 | 182 | 90 | 2 |
| Psf2 (25kDa) | CMG (GINS) | 0 | 145 | 46 | 0 |
| Psf3 (22kDa) | CMG (GINS) | 0 | 60 | 27 | 0 |
| Sld5 (34kDa) | CMG (GINS) | 0 | 358 | 109 | 0 |
| Cdc45 (74kDa) | CMG | 2 | 502 | 300 | 8 |
| Mcm2 (99kDa) | CMG (Mcm2-7) | 18 | 1160 | 735 | 5 |
| Mcm3 (107kDa) | CMG (Mcm2-7) | 5 | 1319 | 790 | 18 |
| Mcm4 (105kDa) | CMG (Mcm2-7) | 9 | 1353 | 924 | 5 |
| Mcm5 (86kDa) | CMG (Mcm2-7) | 11 | 921 | 545 | 16 |
| Mcm6 (113kDa) | CMG (Mcm2-7) | 4 | 1303 | 835 | 20 |
| Mcm7 (95kDa) | CMG (Mcm2-7) | 0 | 1026 | 583 | 13 |
| Ctf4<br>(104kDa) | Replisome | 4 | 1746 | 1275 | 8 |
| Tof1<br>(141kDa) | Replisome<br>(Tof1-Csm3) | 2 | 1015 | 645 | 11 |
| Csm3<br>(36kDa) | Replisome<br>(Tof1-Csm3) | 0 | 132 | 103 | 2 |
| Mrc1<br>(124kDa) | Replisome | 0 | 1280 | 736 | 7 |
| Top1<br>(90kDa) | Replisome | 0 | 329 | 140 | 5 |
| Spt16<br>(119kDa) | Replisome<br>(FACT) | 375 | 1491 | 821 | 260 |
| Pob3<br>(63kDa) | Replisome<br>(FACT) | 95 | 534 | 264 | 88 |
| Pol2<br>(256kDa) | Pol $\epsilon$ | 17 | 210 | 114 | 22 |
| Dpb2<br>(78kDa) | Pol $\epsilon$ | 0 | 38 | 34 | 4 |
| Dpb3<br>(23kDa) | Pol $\epsilon$ | 0 | 10 | 10 | 2 |
| Dpb4<br>(22kDa) | Pol $\epsilon$ | 2 | 19 | 12 | 2 |
| Pol1<br>(167kDa) | Pol $\alpha$ | 5 | 204 | 127 | 19 |
| Pol12<br>(79kDa) | Pol $\alpha$ | 0 | 58 | 57 | 3 |
| Pri1<br>(48kDa) | Pol $\alpha$ | 0 | 44 | 39 | 7 |
| Pri2<br>(62kDa) | Pol $\alpha$ | 0 | 37 | 38 | 4 |
| **Other reported substrates of SCF<sup>Dia2</sup>** |  |  |  |  |  |
| Rad51<br>(51kDa) | - | 2 | 5 | 13 | 7 |
| Cdc6 | ORC-CDC6 | 0 | 0 | 0 | 0 |
| Tec1<br>(55kDa) |  | 0 | 0 | 0 | 3 |
| Sir4<br>(152kDa) |  | 39 | 13 | 45 | 34 |

**Appendix Table S3.**
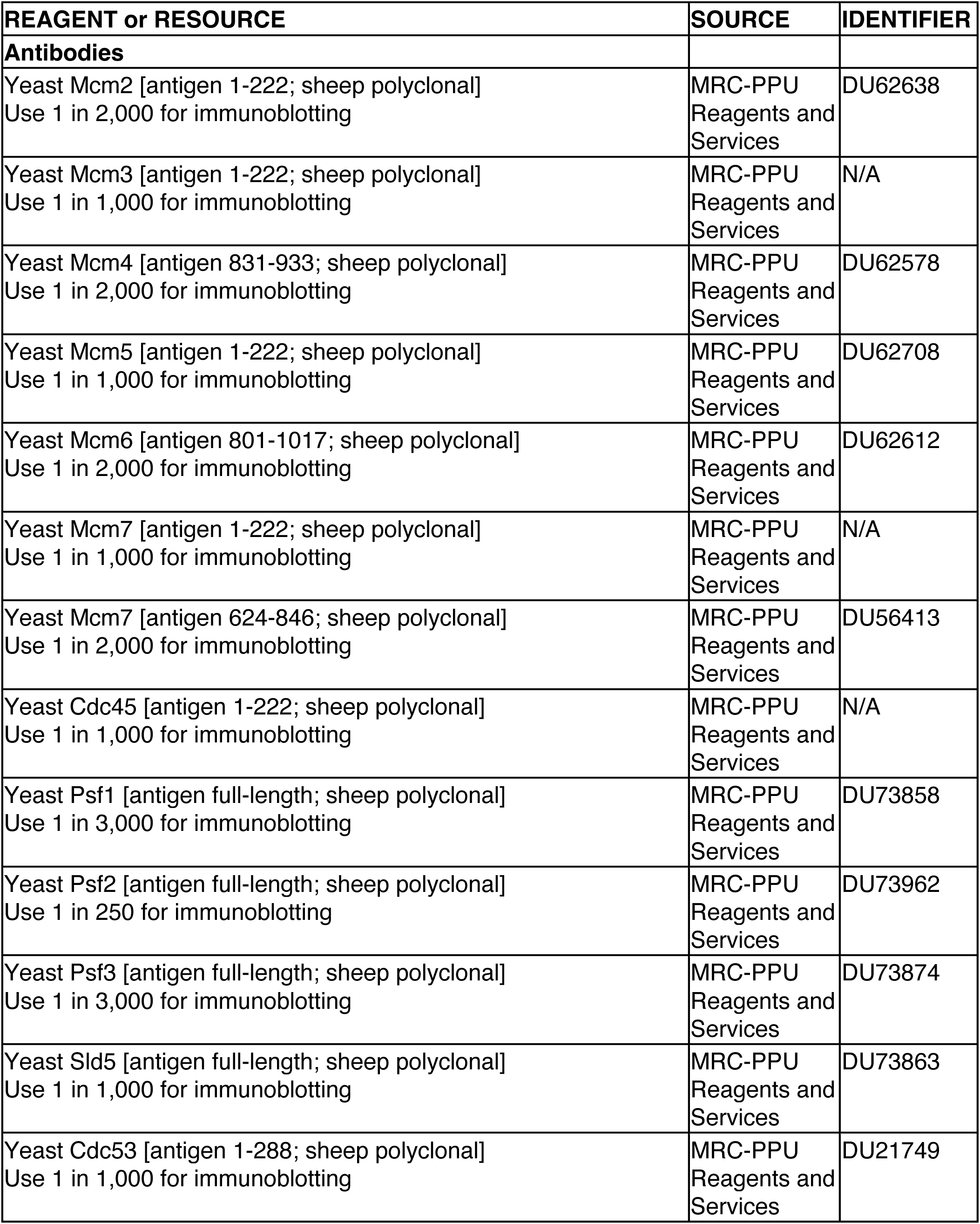
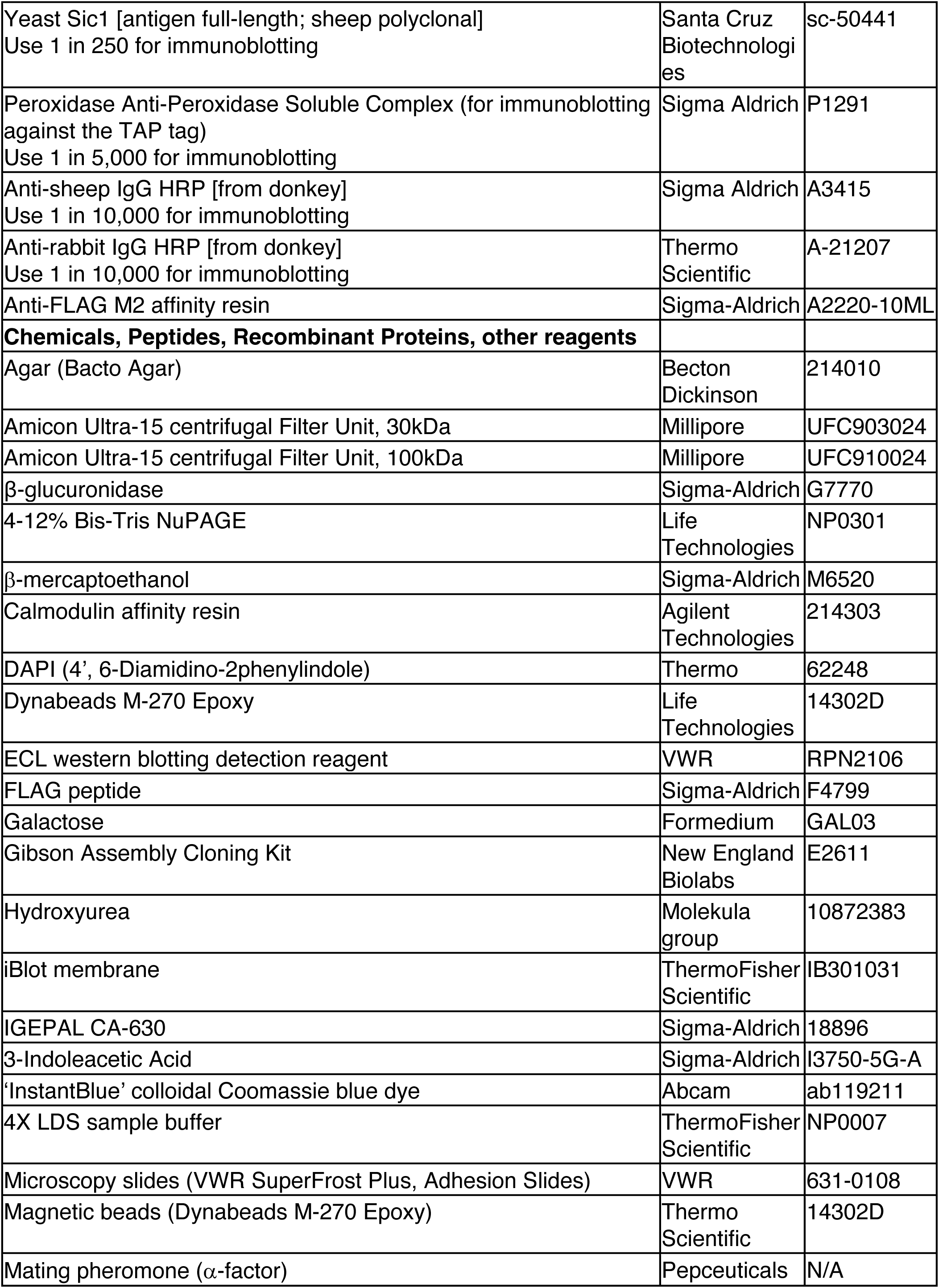
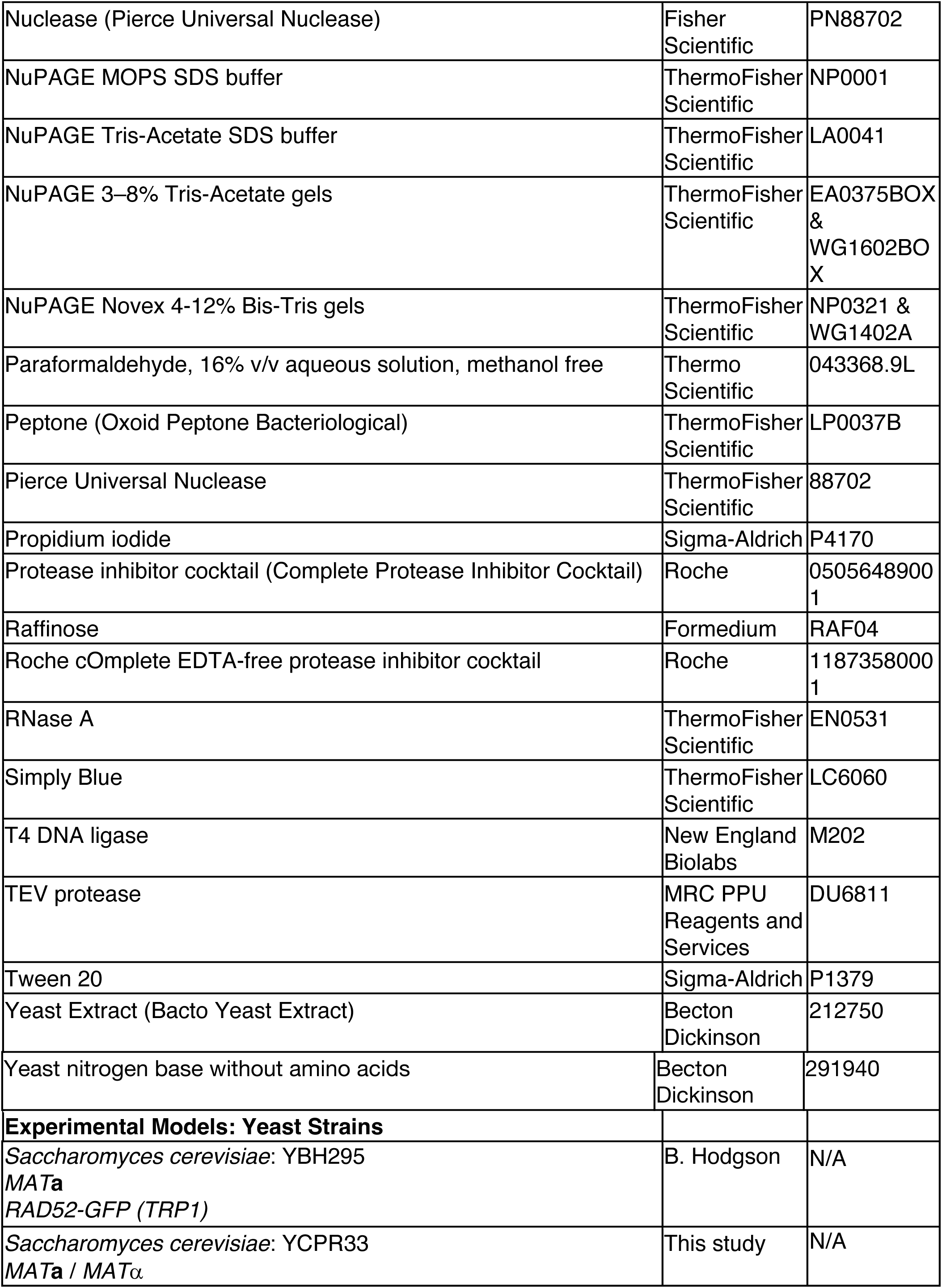
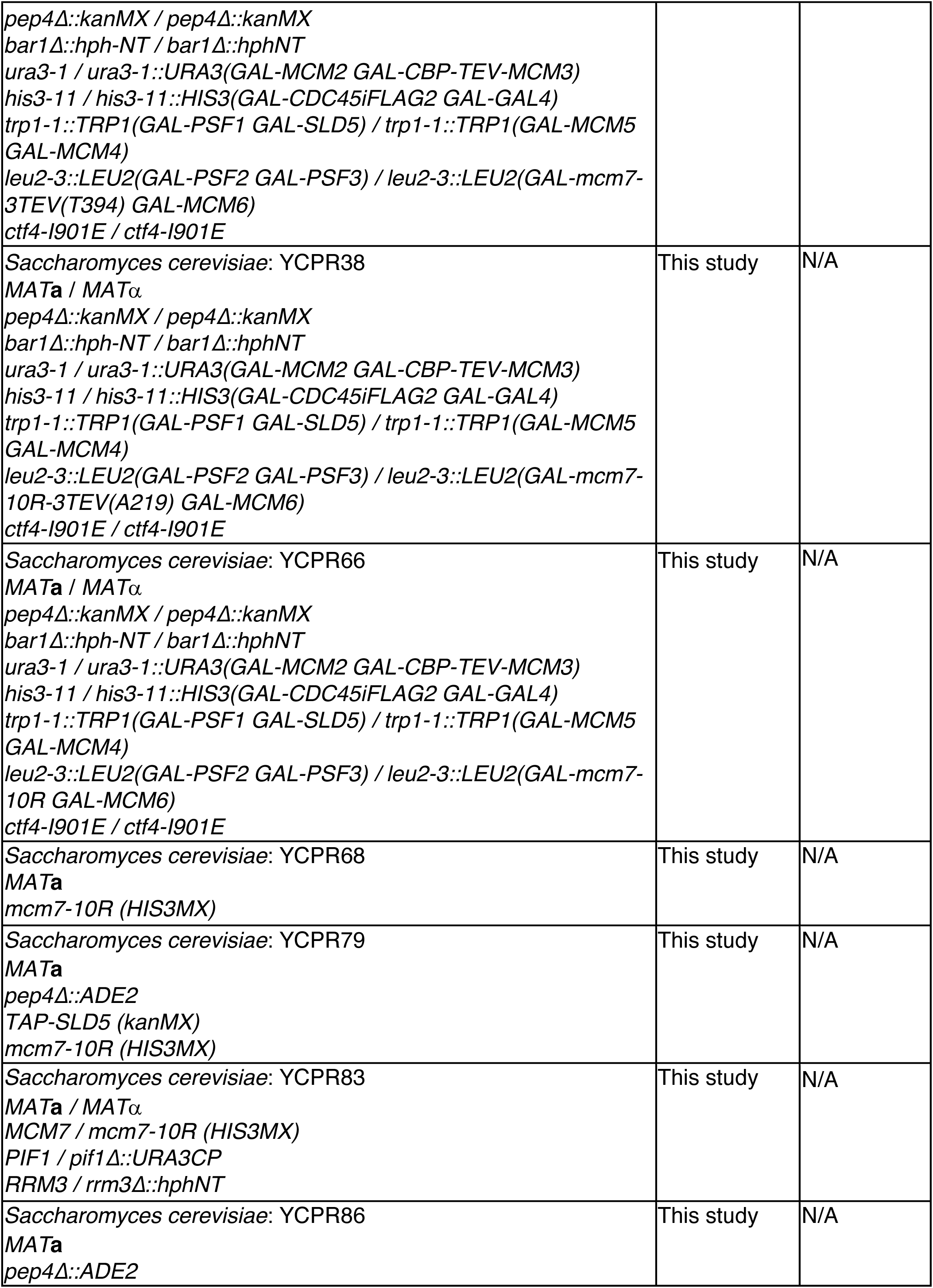
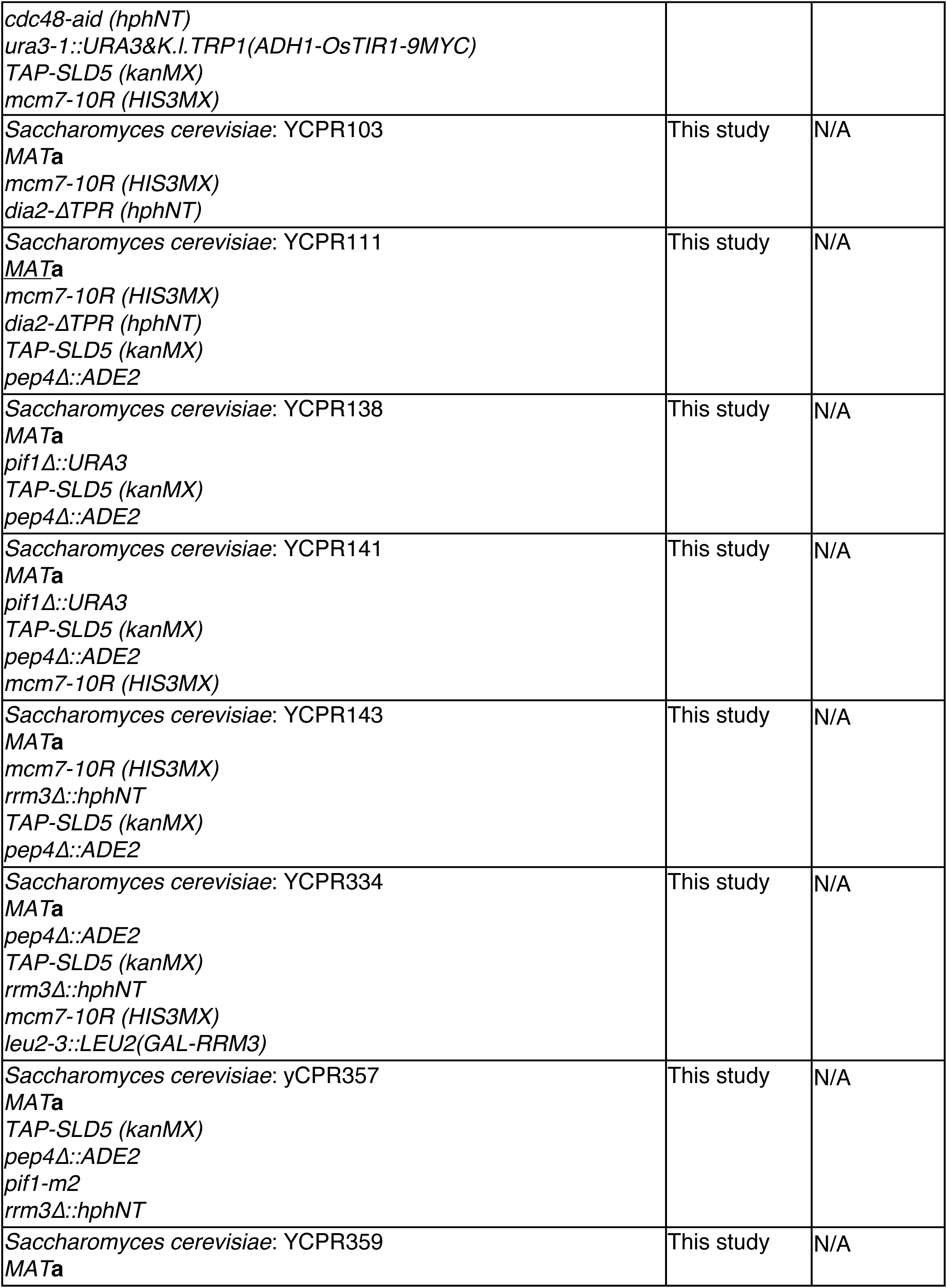
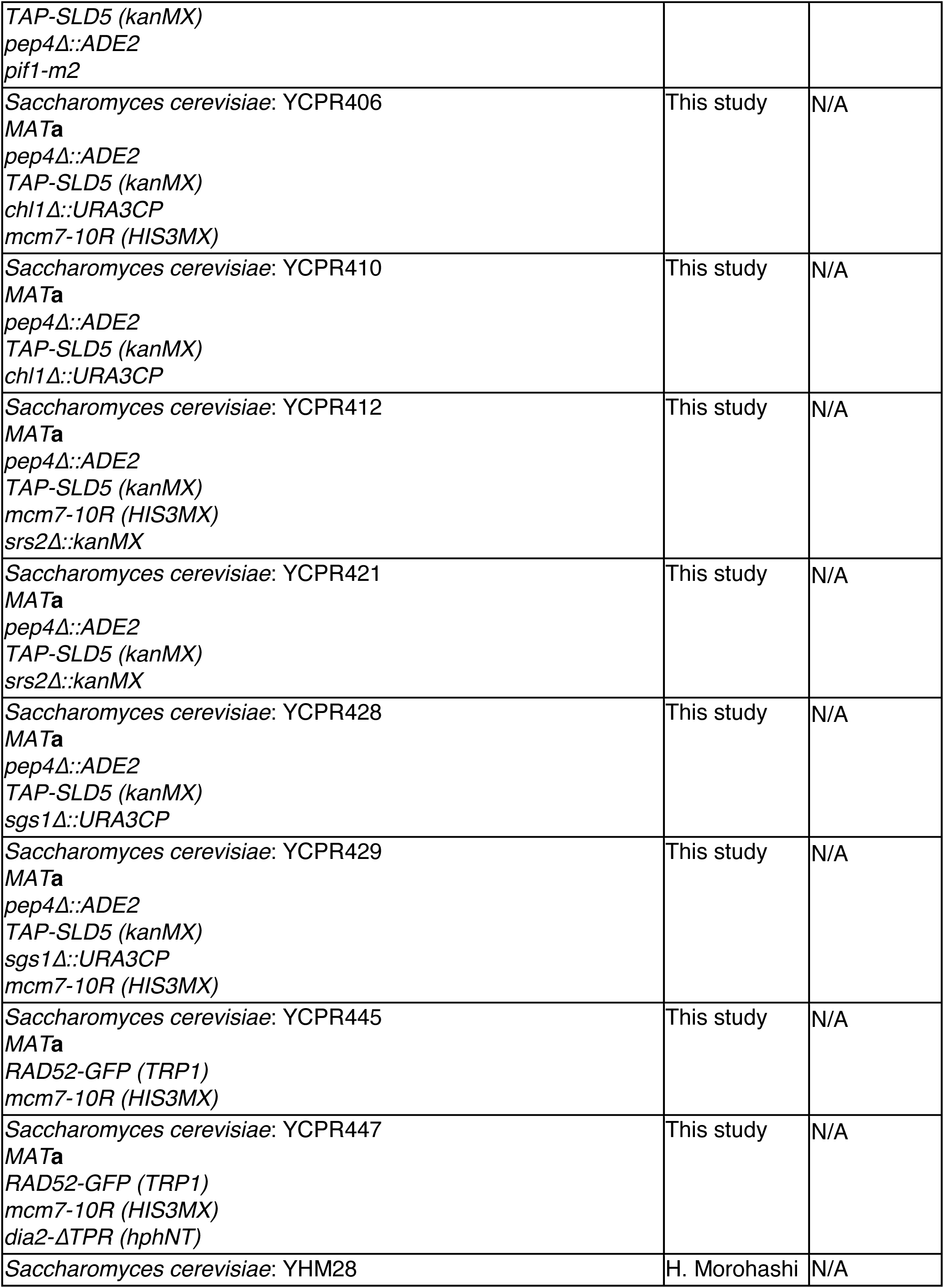
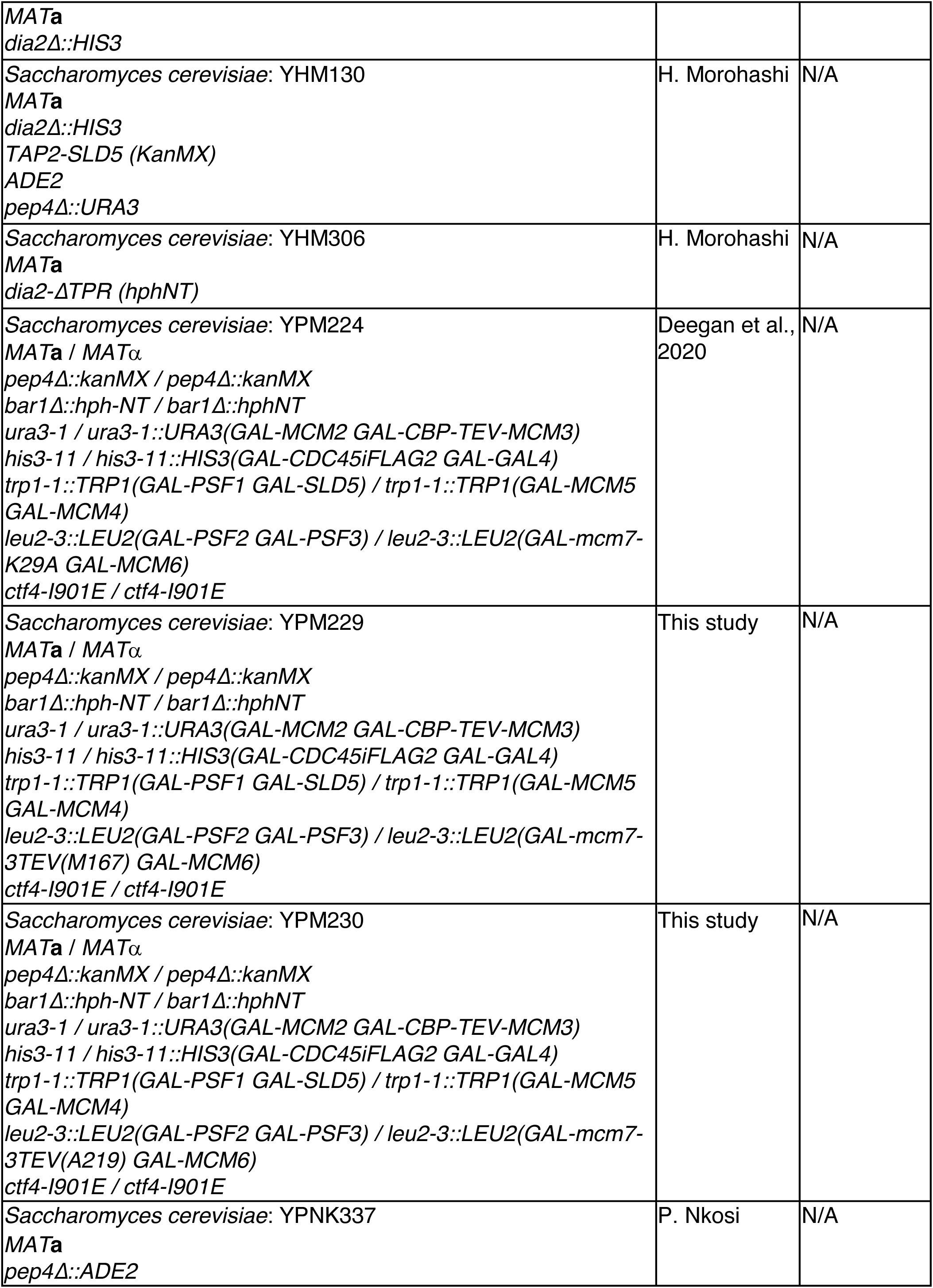
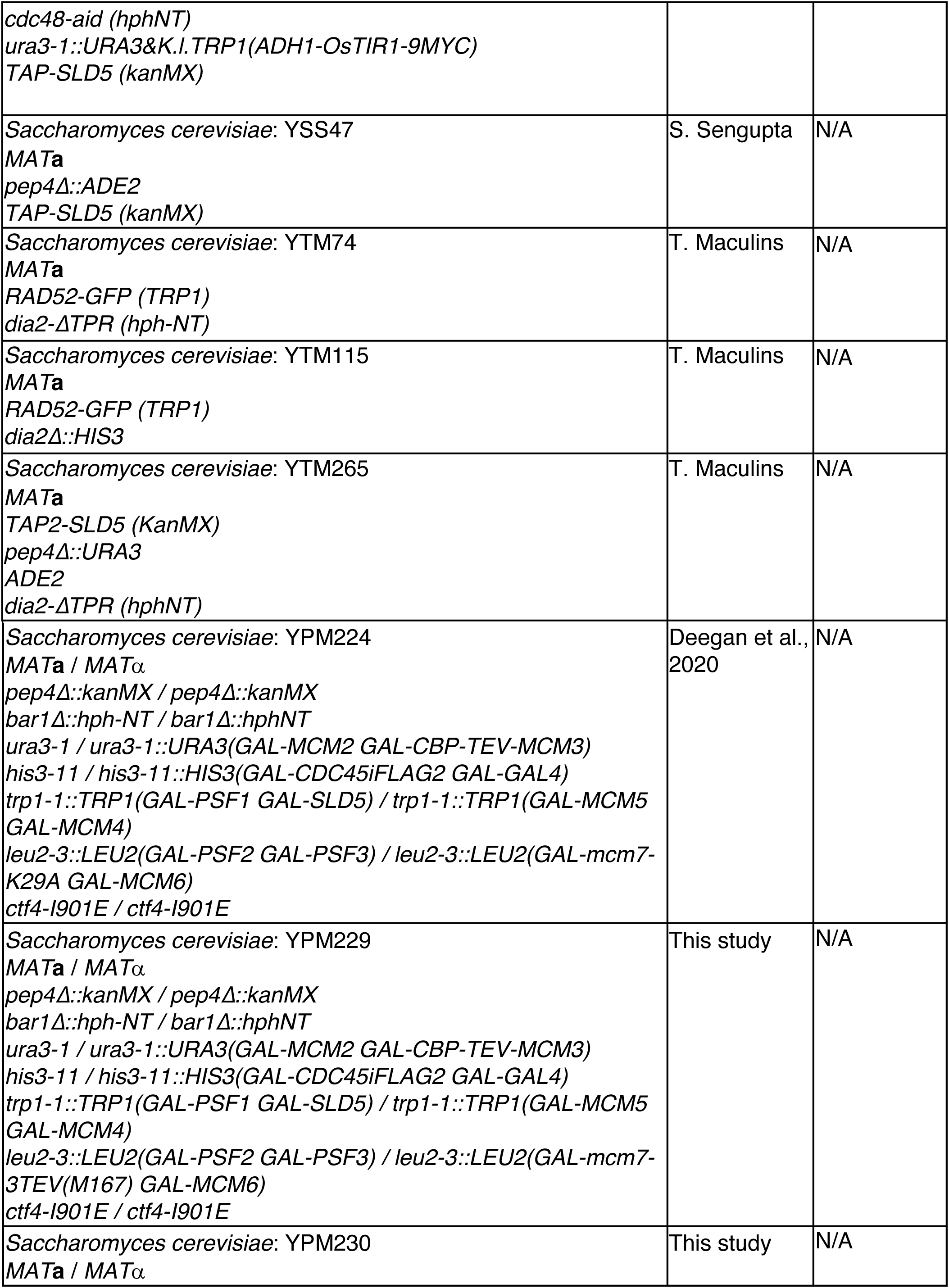
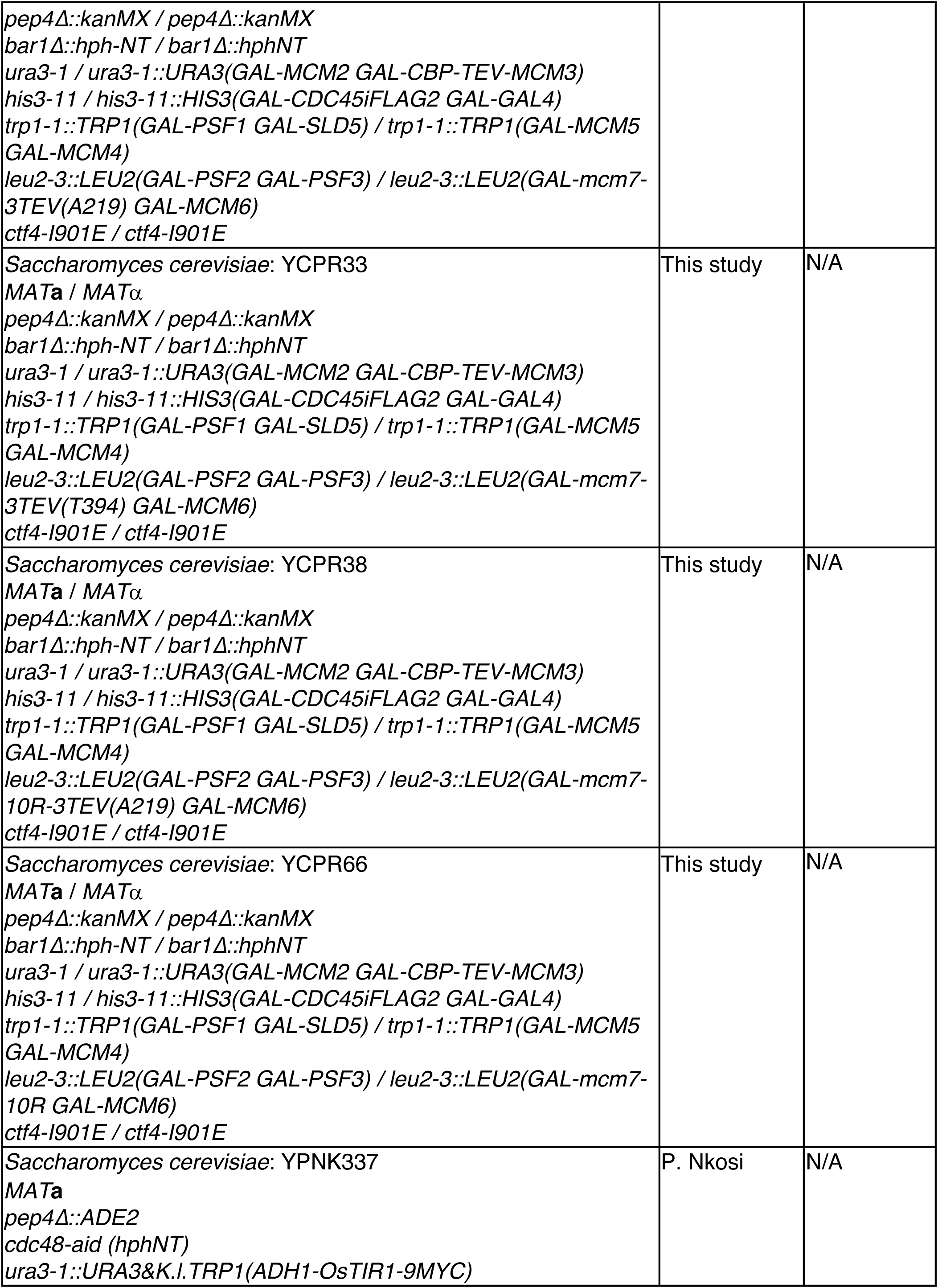
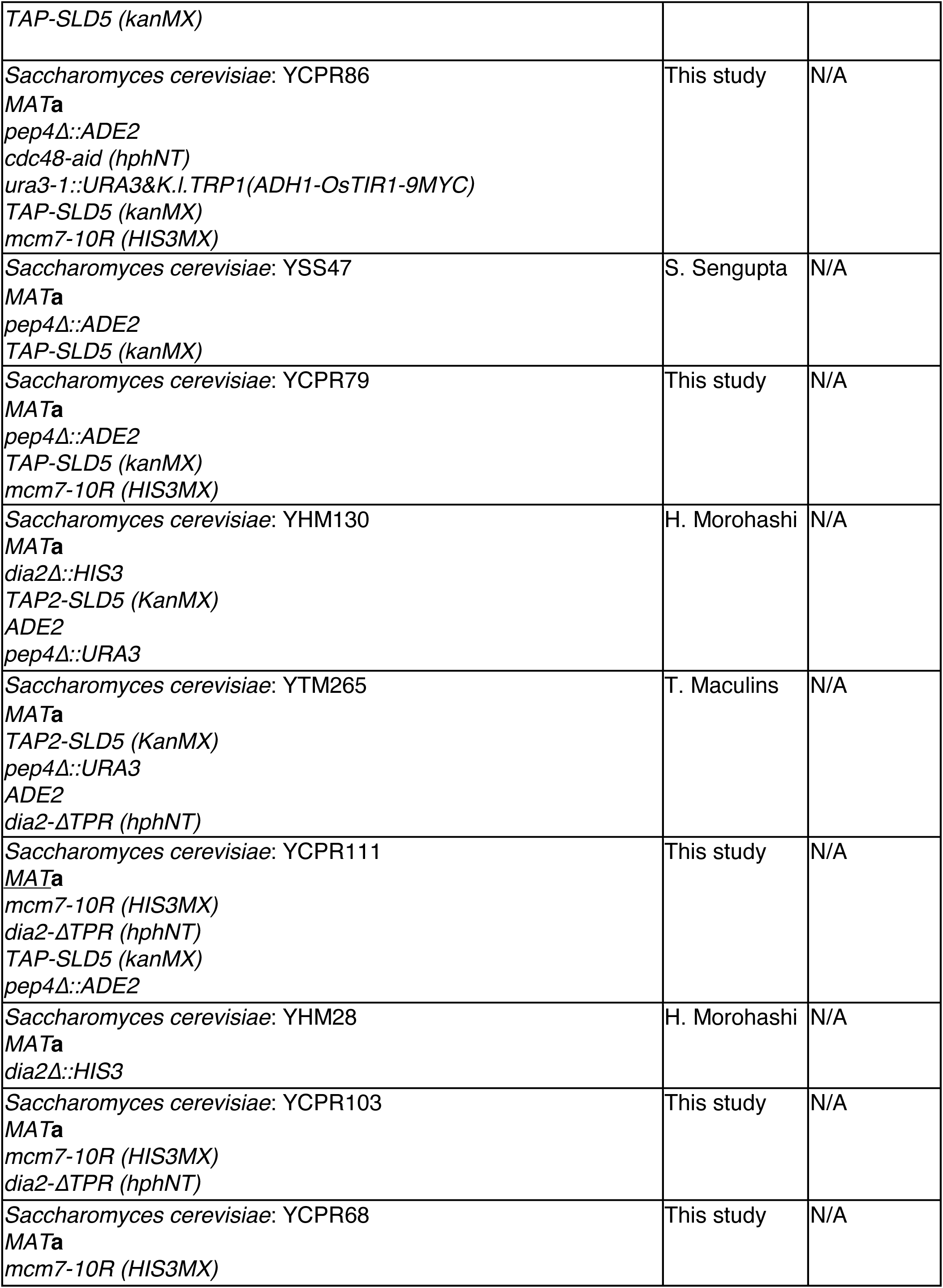
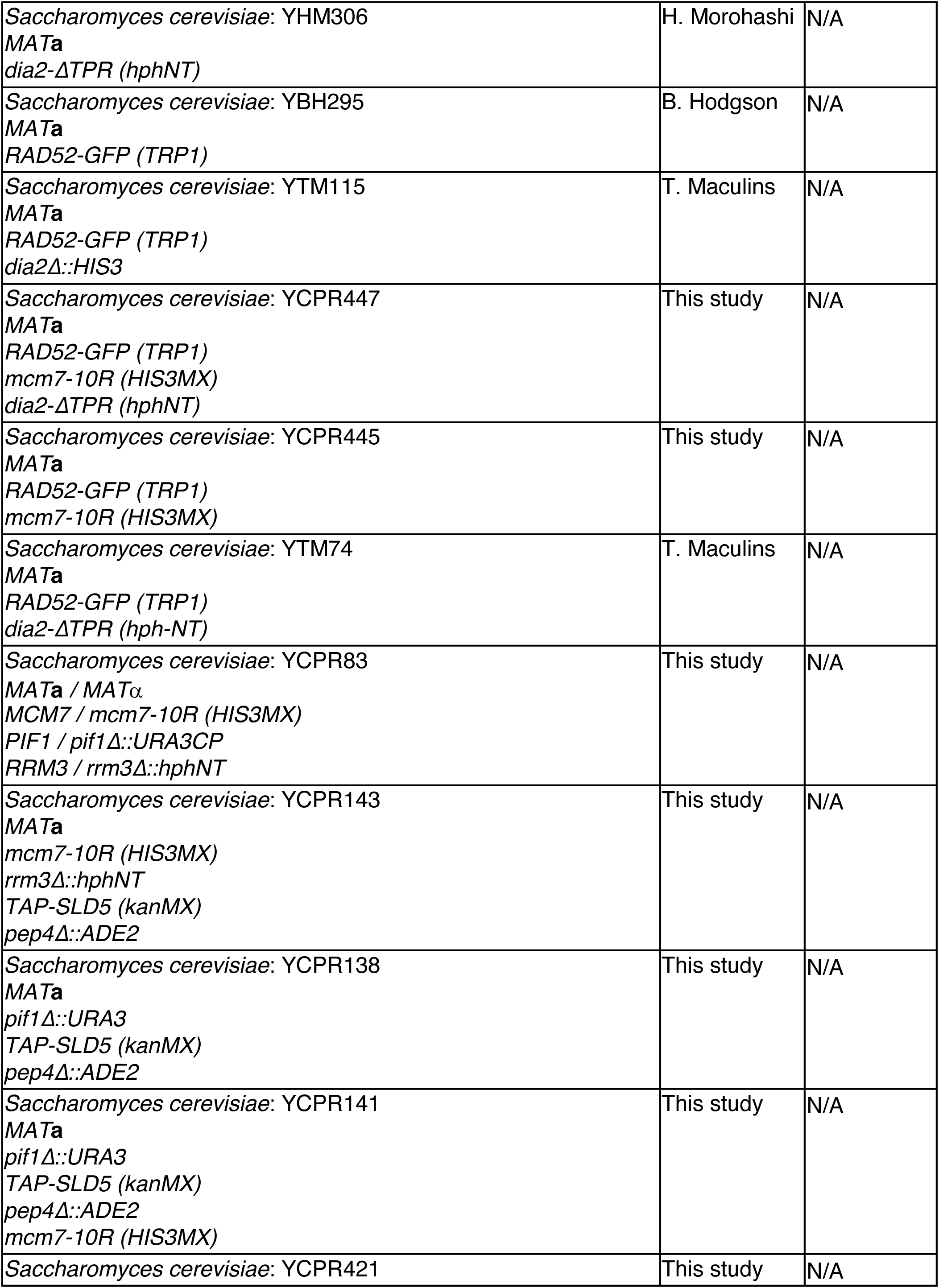
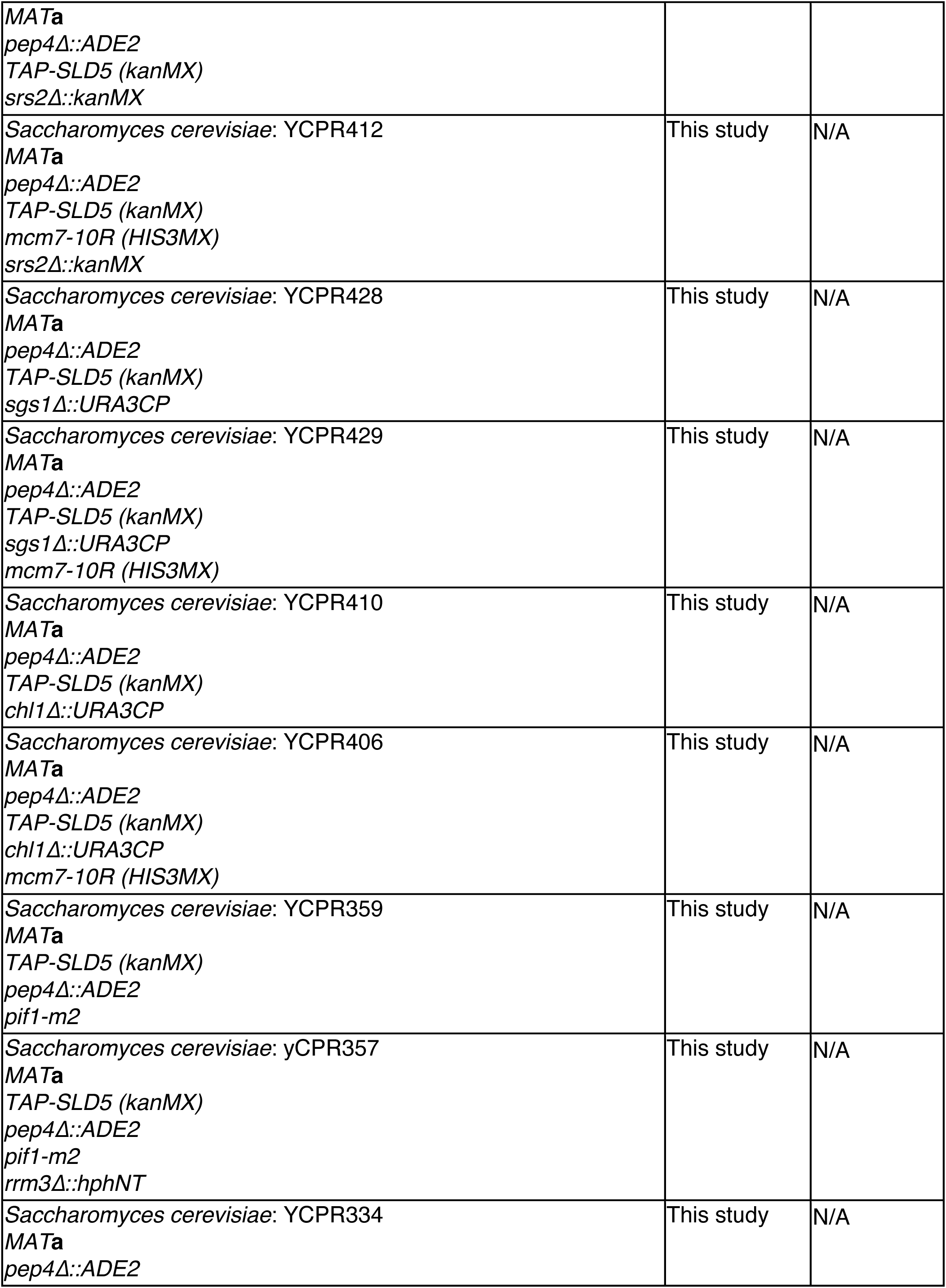
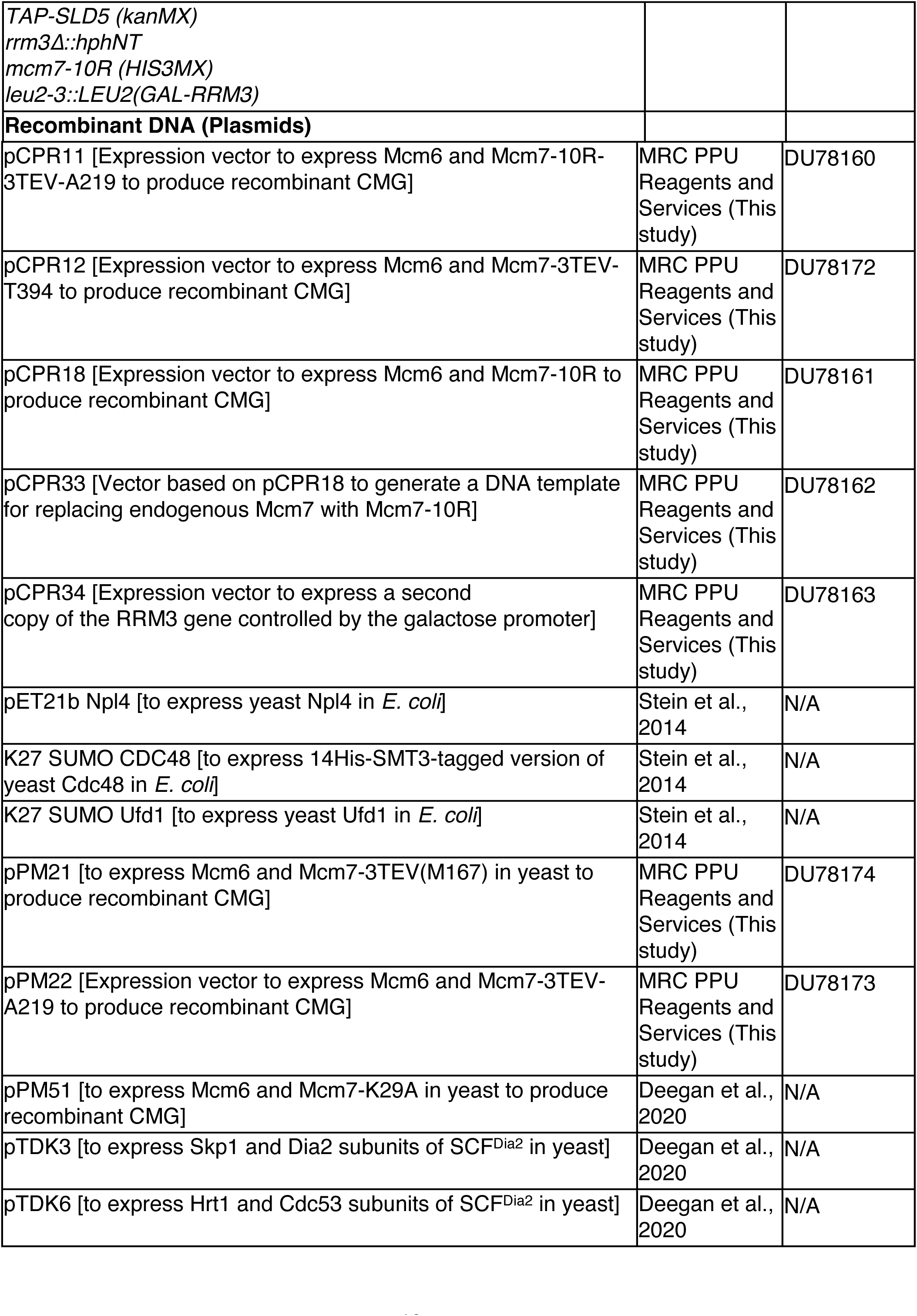
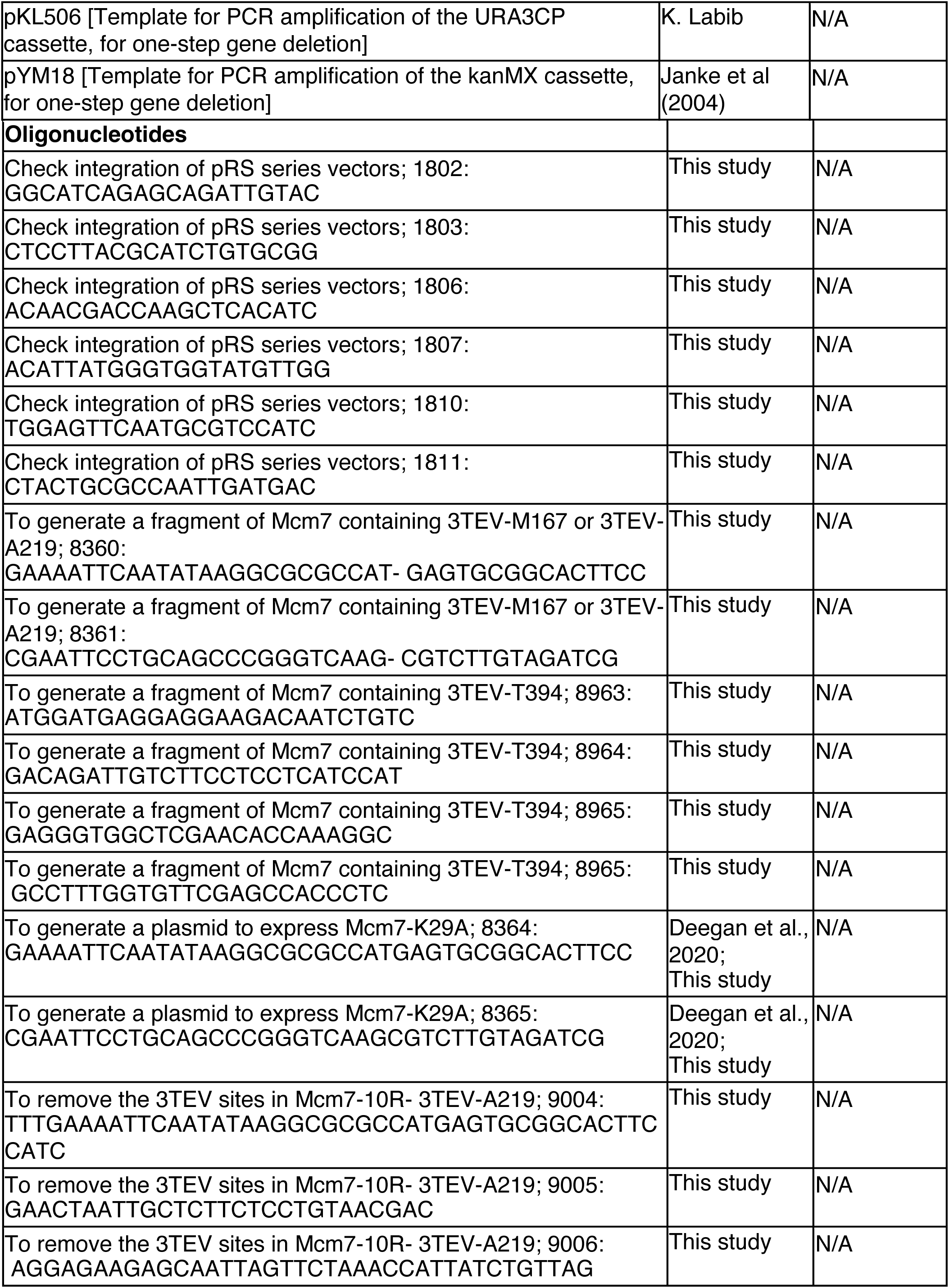
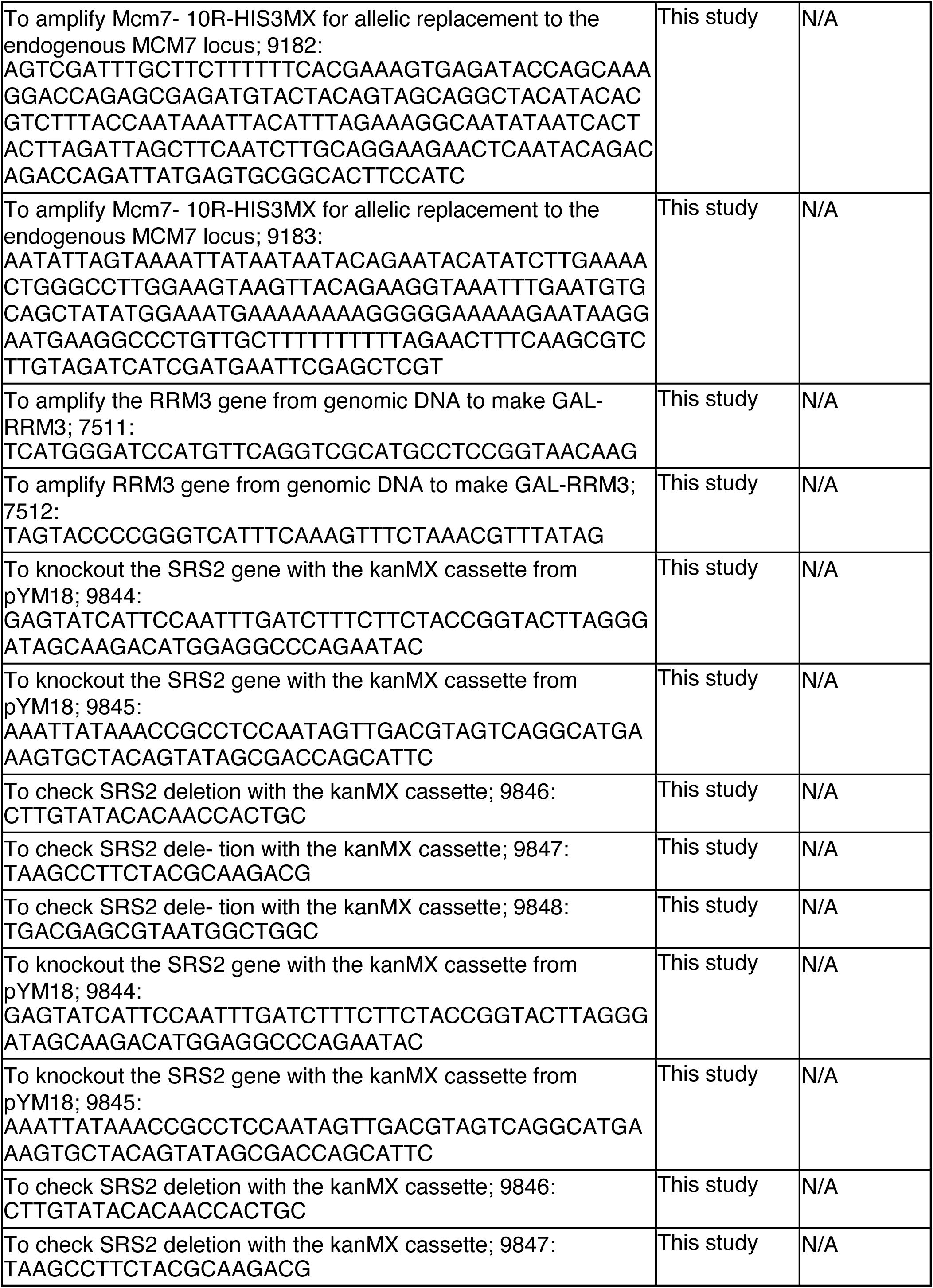
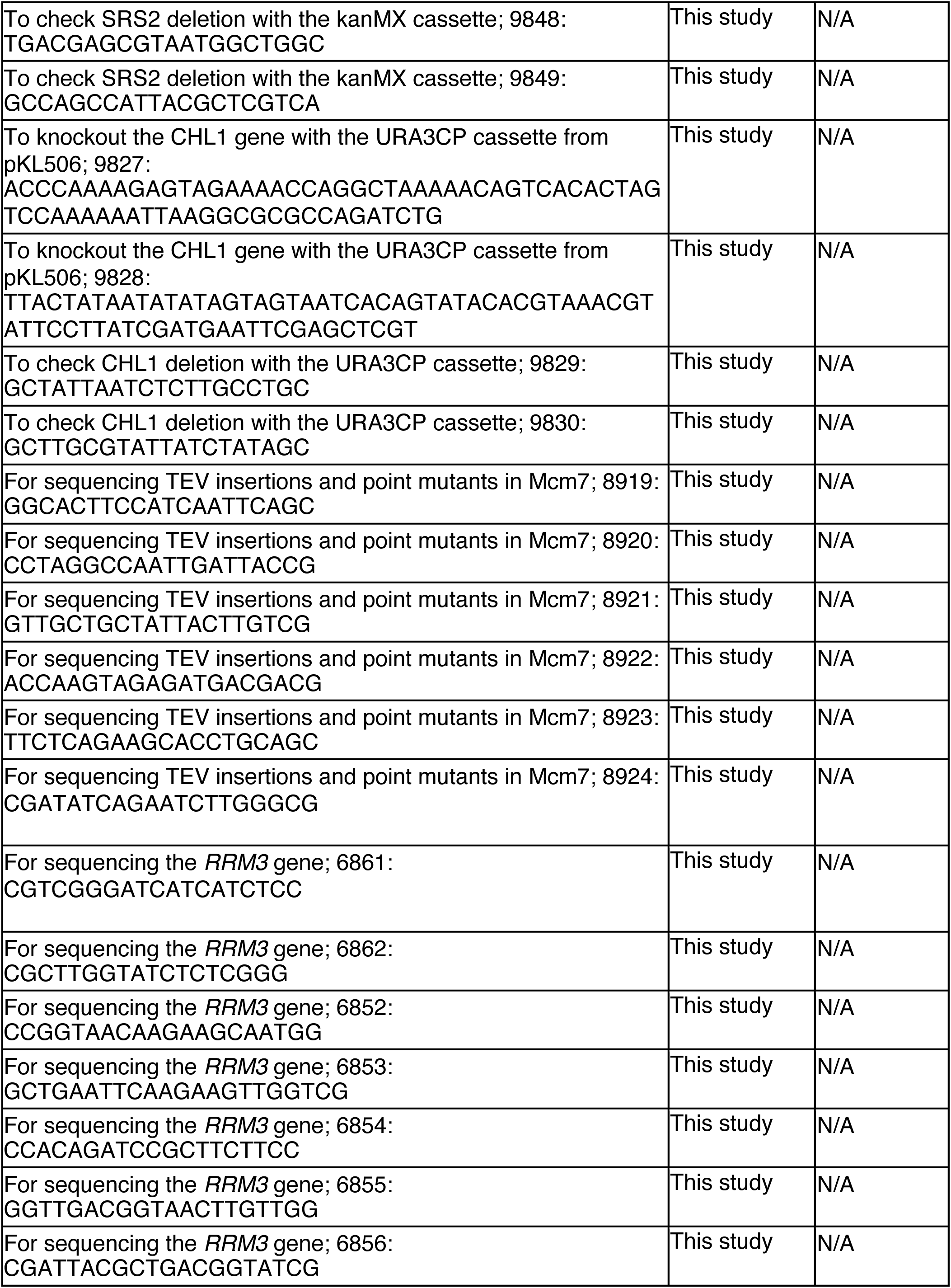
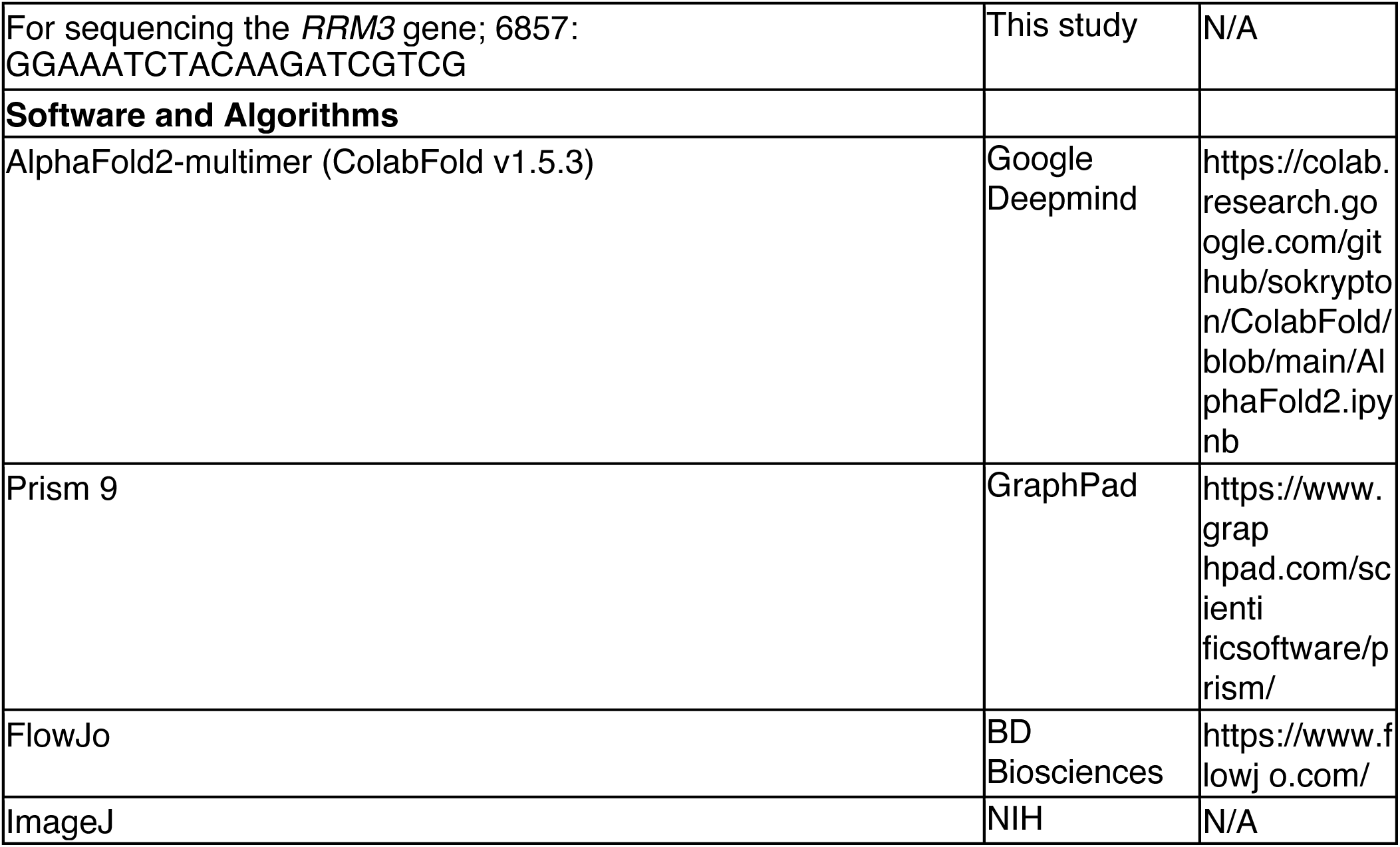
Reagents and resources used in this study.

## Notes

### Competing Interest Statement

The authors have declared no competing interest.

